# Functional drug susceptibility testing based on biophysical measurements predicts patient outcome in glioblastoma patient-derived neurosphere models

**DOI:** 10.1101/2020.08.05.238154

**Authors:** Max A. Stockslager, Seth Malinowski, Mehdi Touat, Jennifer C. Yoon, Jack Geduldig, Mahnoor Mirza, Annette S. Kim, Patrick Y. Wen, Kin-Hoe Chow, Keith L. Ligon, Scott R. Manalis

## Abstract

Functional precision medicine aims to match each cancer patient to the most effective treatment by performing *ex vivo* drug susceptibility testing on the patient’s tumor cells. Despite promising feasibility studies, functional drug susceptibility testing is not yet used in clinical oncology practice to make treatment decisions. Often, functional testing approaches have measured *ex vivo* drug response using metabolic assays such as CellTiter-Glo, which measures ATP as a proxy for numbers of viable cells. As a complement to these existing metabolic drug response assays, we evaluated whether biophysical assays based on cell mass (the suspended microchannel resonator mass assay) or size as measured by microscopy (the IncuCyte assay) could be used as a readout for *ex vivo* drug response. Using these biophysical assays, we profiled the *ex vivo* temozolomide responses of a retrospective cohort of 70 glioblastoma patient-derived neurosphere models with matched clinical outcomes and found that both biophysical assays predicted patients’ overall survival with similar power to MGMT promoter methylation, the clinical gold standard biomarker for predicting temozolomide response in glioblastoma. These findings suggest that biophysical assays could be a useful complement to existing metabolic approaches as “universal biomarkers” to measure sensitivity or resistance to anti-cancer drugs with a wide variety of cytostatic or cytotoxic mechanisms.

**One-sentence summary:** By using biophysical assays to perform *ex vivo* drug susceptibility testing on 70 glioblastoma patient-derived neurosphere models, we find that functional testing predicts the duration that patients survive when treated with temozolomide, the standard of care chemotherapy.

## Introduction

Cancer precision medicine seeks to match each individual patient to the most effective available therapy. To date, precision medicine has largely been based on genomic profiling, in which patients with certain pre-defined genetic alterations are identified and matched with drugs targeting those specific abnormalities. The introduction and rapid success of the first genome-guided drugs in the early 2000s led to a period of optimism regarding the future of genomic precision medicine, with the National Cancer Institute setting a goal of “the elimination of suffering and death due to cancer” by 2015 *(1)*. However, despite this early success, progress in identifying additional actionable mutations has been slow, and today less than 20% of patients with metastatic cancer are eligible for FDA-approved genome-guided drugs *(2)*. This slow progress has led some to speculate that most of the “low-hanging fruit” of actionable mutations have already been identified, and that new, complementary approaches will be needed to continue making progress in improving outcomes for cancer patients *(3)*.

As a complement to genomic precision medicine, functional precision medicine aims to match cancer patients to effective therapies by performing *ex vivo* drug sensitivity testing directly on biopsied tumor cells. The vision of functional precision medicine is to predict drug sensitivity at the level of individual patients by sampling the tumor, exposing the tumor cells to candidate drugs *ex vivo*, then measuring the cellular response using integrative readouts such as cell growth, proliferation, or apoptotic signaling. Functional precision medicine is based on the hypothesis that if a patient’s tumor cells do not change in response to *ex vivo* drug exposure, then the patient would be unlikely to respond to that drug, and vice versa. Unlike genomic precision medicine, which typically focuses on identifying small subsets of patients harboring a few specific, well-understood targetable mutations, functional drug sensitivity testing could potentially be used to test susceptibility to a broad range of drugs regardless of the patient’s specific disease or genomic background. Moreover, functional diagnostic approaches have the advantage of integrating many complex mechanisms determining drug sensitivity versus resistance into a single readout, which could facilitate drug discovery for the remaining majority of cancers currently lacking genome-based predictive biomarkers.

Despite continued efforts from many groups to develop functional assays for drug susceptibility testing in cancer *(4)*, there are still no such assays currently used in clinical practice. A key factor that has limited the translation of functional assays to the clinic is a general lack of studies that directly assess whether *ex vivo* drug susceptibility testing is correlated with measures of patient treatment outcome, such as overall survival duration on therapy. This lack of compelling evidence was the basis for a clinical practice guideline published by the American Society of Clinical Oncology which concluded that, as of 2011, there was not sufficient evidence to support use the use of functional drug susceptibility testing in clinical oncology practice; the group has not published a revised guideline since then *(5)*. Most studies linking functional testing to patient outcome have only performed functional testing on tumor samples with matched patient outcomes for small numbers of patients (*n* = 4-9 patients per study with matched clinical outcomes *(6–9)*). One recent study has shown promise in statistically relevant retrospective patient cohorts: Yin et al. *(10)* used the CellTiter-Glo assay to perform *ex vivo* drug susceptibility testing on “patient-derived tumor-like cell clusters” established from 59 gastric, colorectal, and breast tumors with matched clinical outcomes, and found that the assay achieved good performance in predicting clinical outcomes for a range of drugs.

Assessing the correlation between functional testing and patient outcome is difficult due to the logistical challenges associated with obtaining clinical follow-up from patients after collecting the initial tumor sample. As an alternative to this prospective approach, which was used in the studies described above, here we sought to identify a tractable model system in which to conduct a retrospective study comparing functional drug susceptibility testing with patients’ clinical outcomes. Glioblastoma (GBM) is one cancer in which a large-scale retrospective study is possible: using 3D tumor neurosphere culture techniques, “patient-derived neurosphere models” can be established from primary tumor resections, then stored long-term as viable cells. Patient-derived neurosphere models have been robustly validated to preserve key phenotypic and genotypic features of the patient’s tumor *in vitro*.

In this study, we utilized a retrospective cohort of 70 genomically-characterized patient-derived neurosphere models (generated by the Dana-Farber Cancer Institute Center for Patient-Derived Models) with matched patient treatment history information, including overall survival duration. We measured the response of these GBM patient-derived neurosphere models to temozolomide (TMZ), an alkylating chemotherapy agent that serves as the standard of care treatment for GBM, and then asked whether functional TMZ susceptibility testing correlated with the duration that patients survived after standard of care treatment including TMZ. This study serves as a useful intermediate step between initial validation experiments (using immortalized cell lines with little clinical relevance) and a full-scale prospective clinical trial (using fresh patient tissue, but with additional technical and logistical challenges), allowing us to evaluate whether a particular functional assay predicts treatment outcome in a more tractable model system.

## Results

### Biophysical assays for functional drug susceptibility testing

Although no functional assays have yet reached clinical implementation, most published approaches have used metabolic readouts to measure the response of cells to *ex vivo* drug exposure. Perhaps the most commonly-used metabolic assay for measuring *ex vivo* drug response is CellTiter-Glo (Promega), a commercially-available assay that measures ATP levels as a proxy for numbers of viable, metabolically-active cells under different conditions of drug exposure *(7, 8, 10, 11)*. Changes in ATP levels reflect cell growth inhibition, and can be used as a readout of *ex vivo* drug response. Other groups have used different metabolic readouts, such as tetrazolium dye-based cell viability assays *(11–13)*, as alternative metabolic readouts for *ex vivo* drug susceptibility testing.

To complement these existing approaches, in this study we evaluated whether biophysical (rather than metabolic) measurements could be used to read out the response of tumor cells to *ex vivo* drug exposure (**Fig. 1A**). By measuring drug-induced changes in tumor cells’ biophysical properties, such as size, mass and growth rate, one could potentially obtain a simpler, more direct readout of tumor growth inhibition than could be obtained via metabolic measurements alone. Further, because biophysical properties are fundamentally linked to cell state, we hypothesized that biophysical measurements could potentially be used as a “universal biomarker” to measure sensitivity or resistance to drugs with a wide variety of cytostatic or cytotoxic mechanisms.

**Figure 1.**
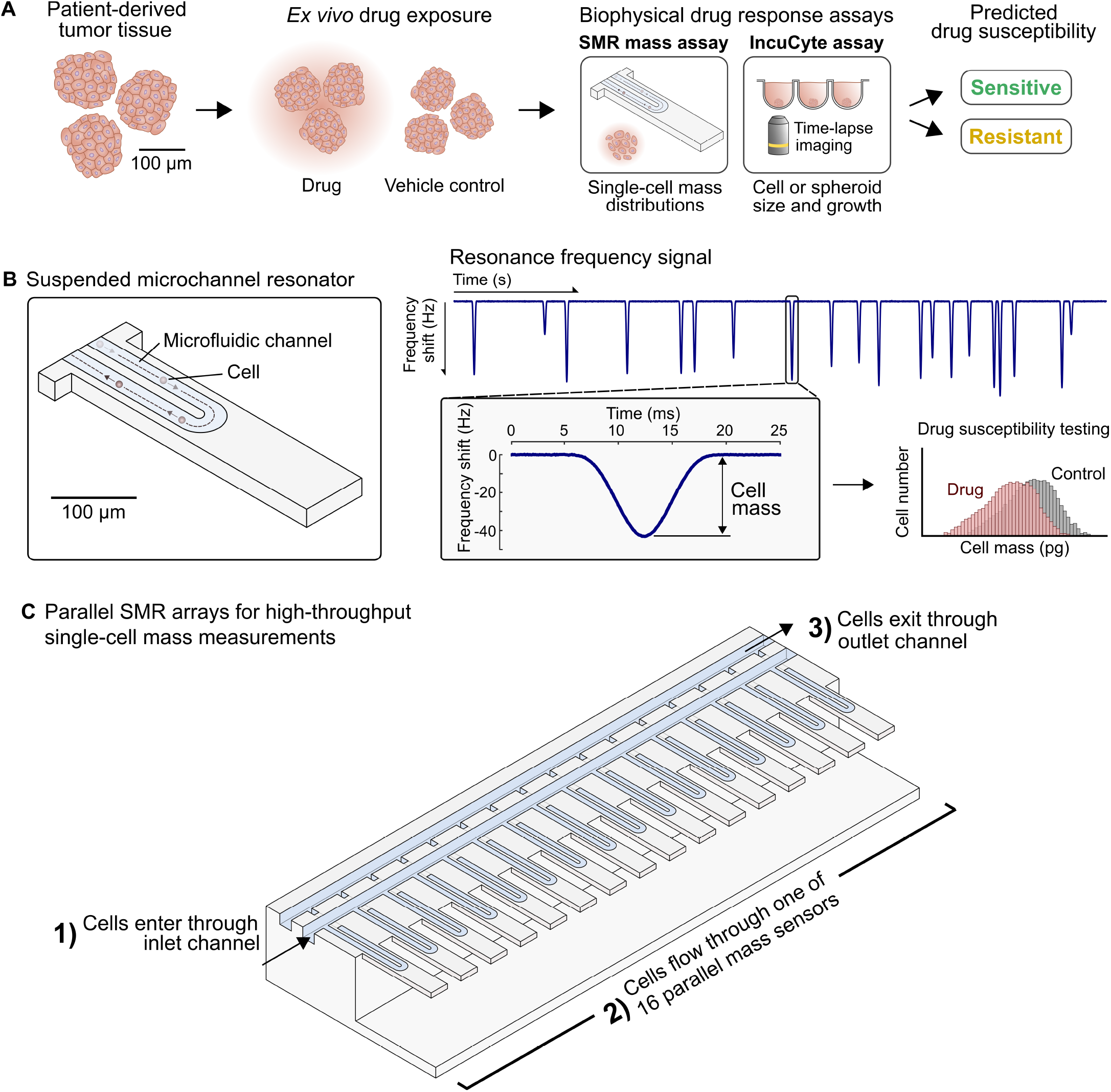
Biophysical assays for functional drug susceptibility testing. **(A)** Workflow for drug susceptibility testing using biophysical assays. Patient-derived tumor tissue is isolated, exposed *ex vivo* to candidate drugs, then biophysical assays are used to measure the response to drug exposure. **(B)** Schematic of a single suspended microchannel resonator (SMR), a microfluidic sensor that weighs single cells as they flow through a resonating micro-cantilever beam. Cell mass is measured by detecting a shift in the cantilever’s resonance frequency as the cell passes through. Cell mass can be used as a readout for *ex vivo* drug susceptibility testing, by comparing the mass of drug-exposed tumor cells to untreated controls. **(C)** Using the parallel SMR array, single-cell mass distributions can be measured at high throughput as cells are simultaneously weighed by multiple SMR sensors in parallel on the same microfluidic chip.

The first biophysical assay, the IncuCyte assay, is based on the IncuCyte S3 Live-Cell Analysis System (Essen Bioscience), a commercially-available microscope optimized for automated time-lapse imaging of cells within a cell culture incubator. Using these images, one can track changes in the growth of individual GBM neurospheres in response to drug exposure as a biophysical readout of drug susceptibility. Versions of the IncuCyte assay have been used predominantly for measuring drug response in adherent 2D cultures *(14)* and to a lesser degree in 3D spheroid cultures *(15)* but without validation in larger scale clinical cohorts.

The second biophysical assay, the suspended microchannel resonator (SMR) assay, is based on a microfluidic sensor that measures buoyant mass (referred to hereafter simply as mass) of single cells by detecting a shift in the resonance frequency of a hollow micro-cantilever beam as cells flow through it (**Fig. 1B**). Using these devices, individual cells can be weighed with a resolution near 50 fg *(16, 17)*, which is highly precise given that the average buoyant mass of cells in this study ranges from 25-100 pg. In prior work, we used a serial array of SMRs to weigh the same cell multiple times over a period of 10-20 minutes to measure mass accumulation rate (MAR), a measure of a single cell’s instantaneous growth rate *(18)*. Changes in MAR can be used as a readout of drug response in tumor cells, even in non-proliferative cells such as multiple myeloma primary tumor samples *(19)*. However, the MAR assay requires several hours to collect growth measurements for each drug condition, thereby limiting its potential clinical implementation.

To address the limited throughput of the MAR assay, we eliminated the need to measure the growth rate of each individual cell, and instead detect drug-induced changes in cell growth by comparing the mass distributions of drug-exposed versus control cell populations (**Fig. 1B**). Recently, high-throughput single-cell mass measurements have been enabled by the parallel SMR array (**Fig. 1C**), a microfluidic device containing sixteen SMRs connected fluidically in parallel and operated simultaneously on the same microfluidic chip *(20)*. By operating many sensors simultaneously, a population of cells can be weighed with maximum throughput of thousands of cells per minute. By measuring the mass of ~2000 cells per sample, changes in mean cell mass as small as 3% can be detected with high statistical power and confidence while using as little as 2 minutes of instrument time per sample. This represents a ~30-fold improvement in the rate at which samples can be measured compared to the MAR assay (**Supplemental Note 1**)

### Validation of the SMR mass assay using conventional cancer cell lines

Unlike the IncuCyte assay, the SMR mass assay has not been previously applied to *ex vivo* drug susceptibility testing. Therefore, as a first step, we used conventional cancer cell lines as a model system to validate that single-cell mass measurements were consistent with expected patterns of drug susceptibility and resistance. We used two model systems: (1) BCR-ABL-positive leukemia cell lines treated with BCR-ABL inhibitors, and (2) EGFR-mutant lung adenocarcinoma cell lines treated with EGFR inhibitors.

First, we replicated a previous experiment in which leukemia cell lines expressing the oncogenic BCR-ABL fusion protein were exposed to the BCR-ABL inhibitors imatinib and ponatinib (**Fig. 2A-D**). We have observed previously that in the cell line Ba/F3 BCR-ABL, exposure to BCR-ABL inhibitors arrests cells in G1 and drastically alters their growth rate distributions *(19)*. Consistent with this finding, the SMR detected a significant reduction in mean cell mass within 8 hours of exposure to 1.4 μM imatinib and 100 nM ponatinib (**Fig. 2A**). For Ba/F3 BCR-ABL T315I, a cell line engineered to express an imatinib-resistant mutant BCR-ABL, cell mass was not significantly reduced in response to 1.4 μM imatinib, but as expected, was reduced in response to 100 nM ponatinib (**Fig. 2B**).

**Figure 2.**
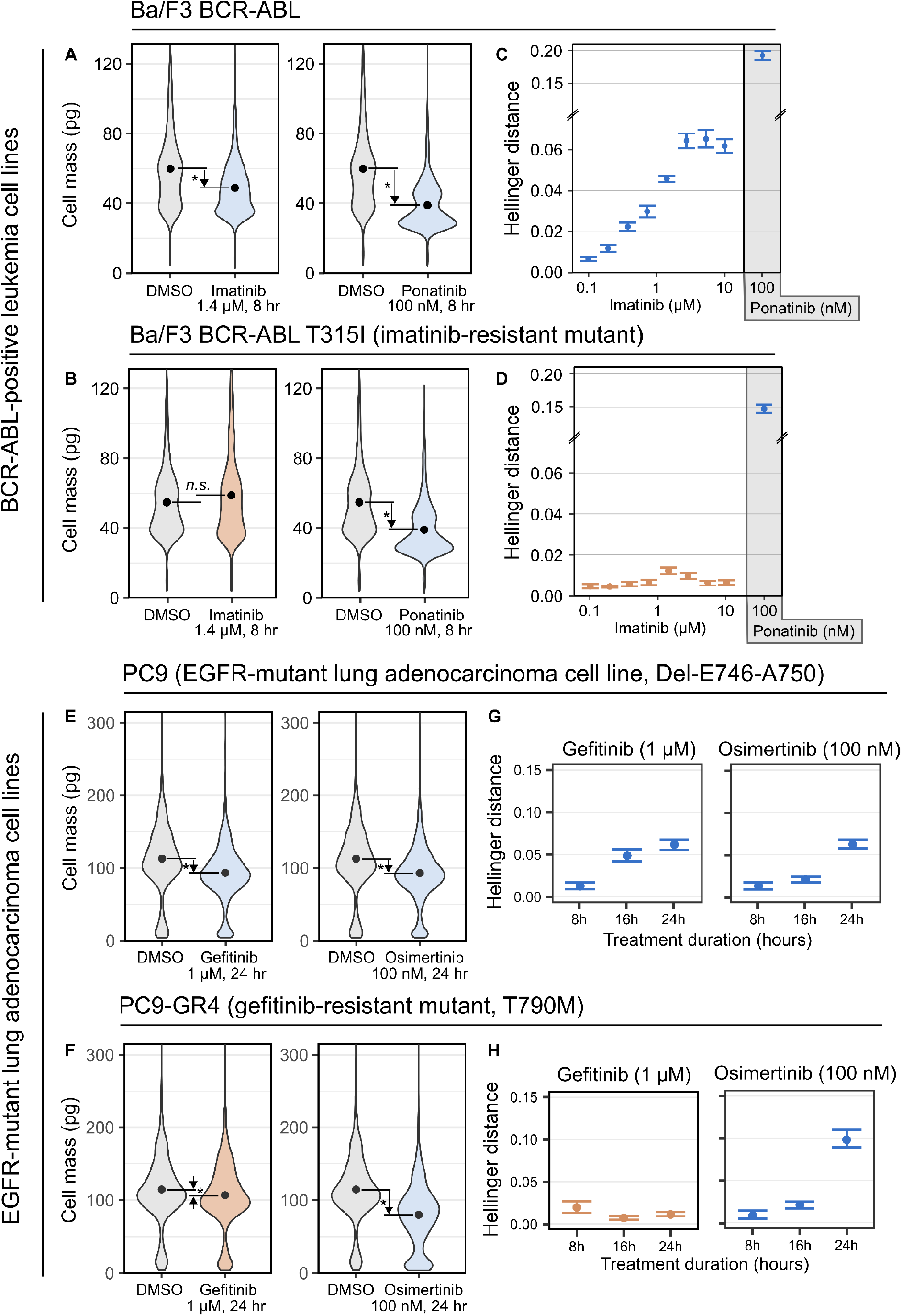
The SMR mass assay recapitulates expected patterns of drug sensitivity and resistance in BCR-ABL-positive leukemia and EGFR-mutant lung cancer cell lines. **(A)** Imatinib and ponatinib drug responses observed in Ba/F3 BCR-ABL. For Ba/F3 BCR-ABL, there is a significant reduction in mean cell mass after 8 hours exposure to 1.4 μM imatinib (one-sided Wilcoxon rank-sum *p* < 10^−3^) and to 100 nM ponatinib (one-sided Wilcoxon rank-sum *p* < 10^−3^). Minimum 2200 cells measured per condition. **(B)** Imatinib and ponatinib drug responses observed in Ba/F3 BCR-ABL T315I. As expected, for Ba/F3 BCR-ABL T315I, there is only a significant reduction in mean mass after 8 hours exposure to 100 nM ponatinib (one-sided Wilcoxon rank-sum *p* < 10^−3^), but not to 1.4 μM imatinib (one-sided Wilcoxon rank-sum *p* > 0.99). Minimum 2200 cells measured per condition. **(C)** Ba/F3 BCR-ABL imatinib dose-response curve measured using the SMR mass assay, given as Hellinger distance ± bootstrap standard error. **(D)** Ba/F3 BCR-ABL T315I imatinib dose-response curve measured using the SMR mass assay, given as Hellinger distance ± bootstrap standard error. **(E)** Gefitinib and osimertinib drug responses observed in PC9, an EGFR-mutant (Del-E746-A750) lung adenocarcinoma cell line expected to be sensitive to both EGFR inhibitors. For PC9, there is a large reduction in mean cell mass in response to 24 hours exposure to either 1 μM gefitinib (16% reduction, one-sided Wilcoxon rank-sum *p* < 10^−3^) or to 100 nM osimertinib (16% reduction; one-sided Wilcoxon rank-sum *p* < 10^−3^). Minimum 3000 cells measured per condition. **(F)** Gefitinib and osimertinib drug responses observed in PC9-GR4, expected to be resistant to gefitinib but not to the second-generation EGFR inhibitor osimertinib. For PC9-GR4, there is only a small reduction in cell mass in response to 24 hours exposure to 1 μM gefitinib (6% reduction, one-sided Wilcoxon rank-sum *p* < 10^−3^), but a larger reduction in response to 100 nM osimertinib (34% reduction; one-sided Wilcoxon rank-sum *p* < 10^−3^). Minimum 3000 cells measured per condition. **(G)** Time response for the PC9 cell line after exposure to EGFR inhibitors, given as Hellinger distance ± bootstrap standard error. **(H)** Time response for the PC9-GR4 cell lines after exposure to EGFR inhibitors, given as Hellinger distance ± bootstrap standard error.

To exploit the increased throughput of the SMR mass assay, we next measured dose-response curves for the Ba/F3 and Ba/F3 BCR-ABL T315I cell lines after 8 hours exposure to imatinib (**Fig. 2C-D**). Instead of comparing the mean mass between drug-exposed cells and untreated controls, we computed an alternative summary statistic, the Hellinger distance, to evaluate to what extent the cell mass distributions were altered by drug exposure. The Hellinger distance is a statistic that measures the degree of difference between the mass distributions of drug-treated and untreated cells, where larger Hellinger distances reflect greater differences between the treated and untreated mass distributions (**Methods**). A drug treatment with no effect on the mass distribution would have a Hellinger distance of zero, while drug treatments causing larger shifts in the cell mass distribution would be assigned larger Hellinger distances, up to a maximum of one. The Hellinger distance statistic has the advantage of capturing effects other than changes in cell mass, such as the accumulation of small debris in the culture due to drug-induced cell death, or the emergence of cell subpopulations of different sizes in response to drug exposure. As expected, Hellinger distance increased with imatinib dose for Ba/F3 BCR-ABL but not for Ba/F3 BCR-ABL T315I (**Fig. 2C-D**).

Next, we asked whether the SMR mass assay could also detect expected patterns of drug sensitivity and resistance in a solid tumor cell line model. We used PC9, an EGFR-mutant human lung adenocarcinoma cell line (EGFR Del-E745-A750) known to be sensitive to the EGFR inhibitor gefitinib. Additionally, we used PC9-GR4, a resistance model containing EGFR T790M, a secondary mutation which is known to confer resistance to gefitinib *(21)*. As expected, 1 μM gefitinib induced a large reduction in population cell mass in PC9 after 24 hours of drug exposure (**Fig. 2E**, 19% reduction), but only a small change in PC9-GR4 (**Fig. 2F**, 6% reduction). Also consistently, for both cell lines we observed a large reduction in population cell mass after 24 hour exposure to 100 nM osimertinib, an EGFR inhibitor that also inhibits EGFR T790M *(21)* (16% and 34% mean mass reduction respectively for PC9 and PC9-GR4).

In addition to measuring the cell mass response after 24 hours of drug exposure, we also captured the transient gefitinib and osimertinib responses by sampling the cell mass distributions at 8, 16, and 24 hours of exposure (**Fig. 2G-H**). As expected, the Hellinger distance increased over time for PC9 but not PC9-GR4 when exposed to gefitinib, but increased over time for both cell lines when exposed to osimertinib. Therefore, cell mass measurements reliably and sensitively identified expected patterns of drug sensitivity and resistance in both liquid and solid tumor cell line model systems with vastly increased throughput compared to previous SMR-based approaches.

### Biophysical functional assays for predicting temozolomide sensitivity in glioblastoma

After validating the SMR mass assay in conventional cancer cell lines, we next asked whether functional drug susceptibility testing using the IncuCyte and SMR mass assays could predict the response of glioblastoma patients to temozolomide (TMZ). For comparison, we also measured TMZ response using the CellTiter-Glo metabolic assay due to its frequent use in *ex vivo* drug susceptibility testing *(7, 8, 11)*. Our workflow for functional testing proceeded as follows (**Fig. 3A, Methods**). Neurosphere cultures were dissociated, then seeded as single cells in ultra-low attachment microtiter plates. After allowing 24 hours for cells to re-aggregate into neurospheres, the cells were exposed to either 20 μM temozolomide (a dose comparable to IC_50_ values reported previously *(22)*) or a vehicle control. Neurospheres were monitored for two weeks after TMZ exposure, with functional readouts being taken either continuously (IncuCyte assay) or at fixed timepoints (3, 5, 7, 10, 12, and 14 days of TMZ exposure, for the SMR mass assay and CellTiter-Glo assay) with feeding at regular intervals over the course of drug exposure. For the SMR mass assay, neurospheres were dissociated to single cells prior to measurement by treatment with Accutase.

**Figure 3.**
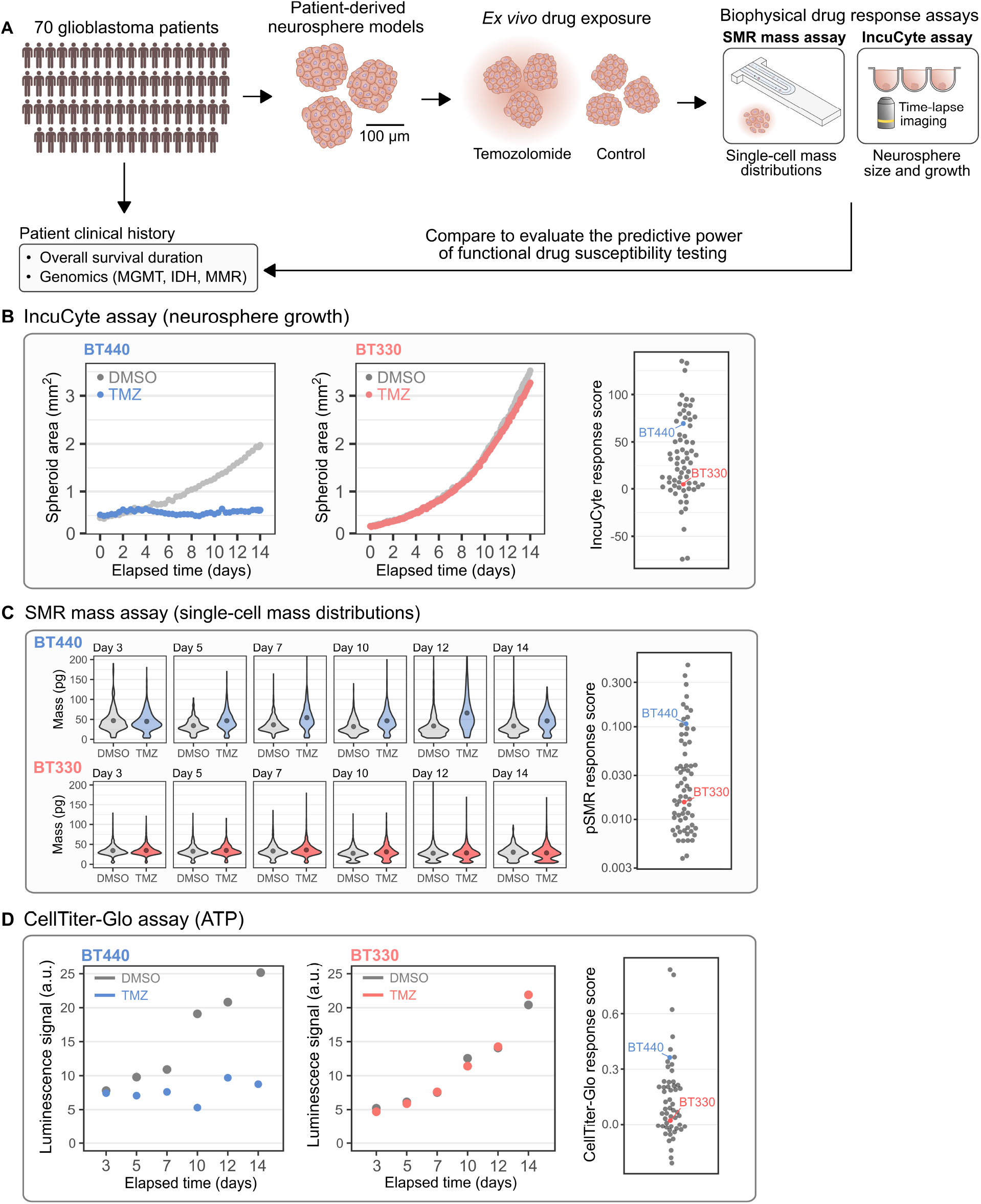
Profiling TMZ responsiveness in glioblastoma patient-derived neurosphere models. **(A)** Workflow for performing functional drug susceptibility testing in glioblastoma (GBM). Patient-derived neurosphere models are established from primary tumors, exposed to temozolomide or a vehicle control, then different biophysical assays are used to measure the TMZ response at multiple timepoints of drug exposure. Biophysical assay results are compared against MGMT promoter methylation status and overall survival duration. **(B)** For the IncuCyte assay, most TMZ-responsive models (e.g., BT440) exhibit a small reduction in spheroid sizes compared to baseline and show marked lack of growth relative to DMSO controls. **(C)** For the SMR mass assay, TMZ-responsive models generally increase in mean cell mass over time compared to controls. **(D)** For the CellTiter-Glo assay, TMZ-responsive models have generally reduced CTG luminescence signals compared to controls.

We performed biophysical functional drug susceptibility testing on a total of 70 patient-derived neurosphere models, which have full clinical annotation and genomic profiling for key biomarkers known to affect TMZ response, including MGMT promoter methylation status (for both the model and the patient), and mismatch-repair (MMR) deficiency status *(23)*. We observed a wide range of biophysical TMZ responses in the patient-derived neurosphere models (**Fig. 3B-D**). In most TMZ-responsive models (e.g., BT440), the IncuCyte assay measured a small reduction in neurosphere size compared to baseline, and a marked reduction in growth relative to DMSO-treated controls (**Fig. 3B**). Interestingly, in TMZ-responsive models the SMR mass assay typically measured an increase in single-cell mass following treatment, consistent with the fact that TMZ arrests sensitive cells at the G2/M checkpoint *(24)*, where individual cells typically have double their initial mass just prior to division (**Fig. 3C**). In TMZ-responsive models, the CellTiter-Glo assay typically measured a reduction in ATP levels relative to controls (**Fig. 3D**). Other patient-derived models (e.g., BT330) showed a striking lack of TMZ response for all three functional assays (**Fig. 3B-D**).

To quantify the TMZ responsiveness of each model, we defined a “response score” for each functional assay, a single statistic describing the extent to which each functional readout changes when the cells are exposed to TMZ (**Fig. 3B-D**, right; **Methods**). For all three assays, a greater response score indicates a larger response to drug exposure. Briefly, the IncuCyte response score is calculated by integrating the difference in spheroid size (as measured by total masked spheroid image area per well) between vehicle and TMZ-exposed conditions over the duration of drug exposure, then normalizing to the growth rate of the vehicle-treated cells; larger IncuCyte response scores reflect greater inhibition of spheroid growth by the drug. Normalizing the response score to the growth rate of DMSO treated cells is critical, since models vary widely in their absolute untreated growth rates, which would otherwise affect the measured degree of growth inhibition as described previously *(25, 26)*. However, normalization corrects for this effect, and after normalization the IncuCyte response score is uncorrelated with the growth rate of untreated cells (Pearson *R* = 0.13; **Supplemental Fig. 1**). The SMR mass response score is defined as the Hellinger distance *(27)* between the vehicle and TMZ-treated cell mass distributions, averaged across the timepoints of drug exposure. The Cell-Titer Glo (CTG) response score is defined as the average of the CTG luminescence signal (normalized to the control at each timepoint) across the timepoints of drug exposure; larger CTG response scores correspond to greater reduction in ATP levels in drug-exposed samples relative to controls. Of note, for all three functional assays, the functional response scores are not clearly bimodal, indicating a continuous spectrum of TMZ responsiveness rather than binary TMZ-sensitive or TMZ-resistant phenotypes.

Of the 70 patient-derived neurosphere models on which any functional testing was performed, data was successfully collected for 68/70 models using the SMR mass assay (full dataset shown in **Fig. 4**), 68/70 models using the IncuCyte assay (full dataset shown in **Supplemental Fig. 2**), and 55/70 models using the CellTiter-Glo assay (full dataset shown in **Supplemental Fig. 3**).

**Figure 4.**
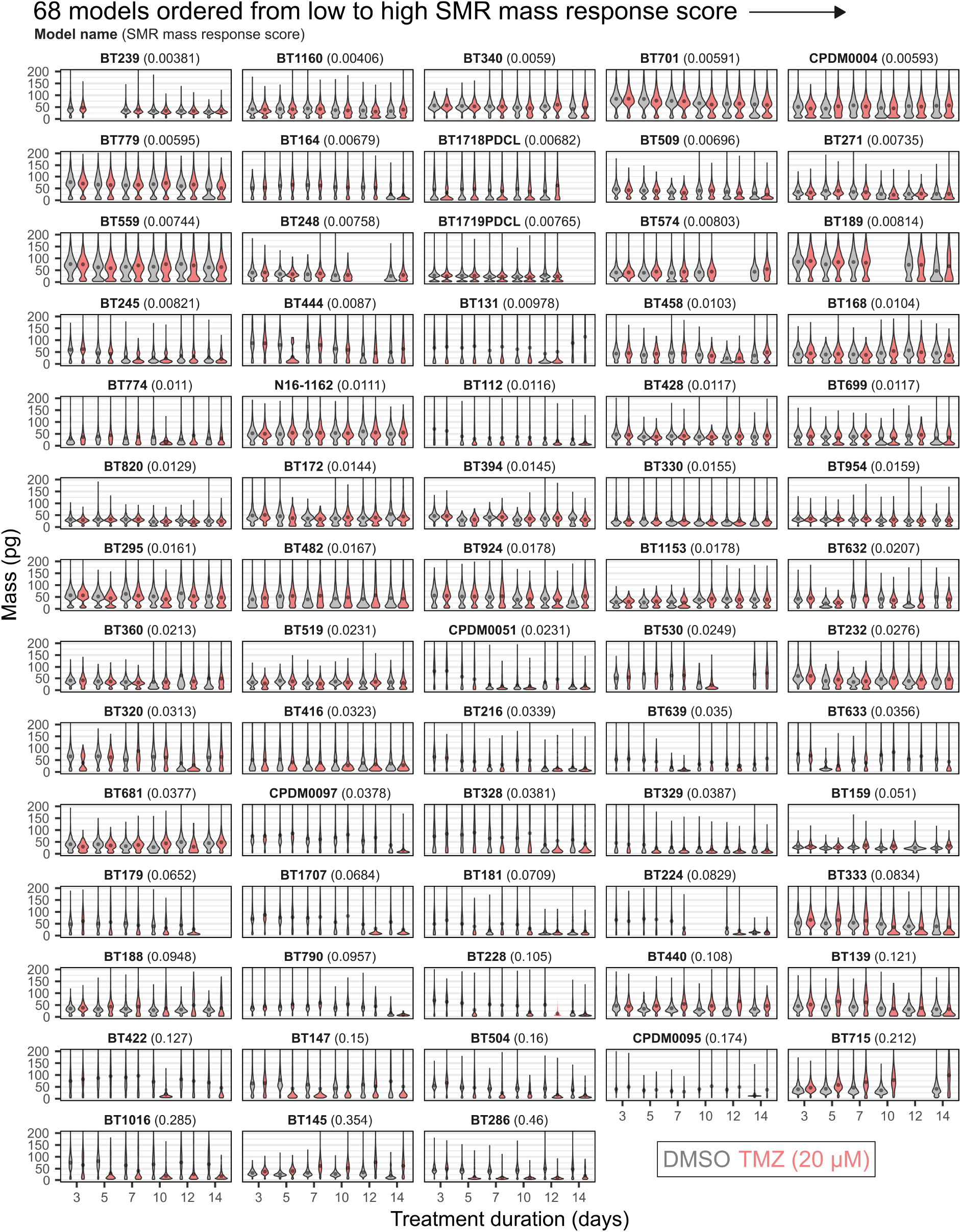
Single-cell mass measurements (SMR mass assay) for profiling the TMZ response of 68 GBM patient-derived neurosphere models. Models are ranked in order of low to high SMR mass response score, which is based on the Hellinger distance between the mass distributions of DMSO and TMZ-treated cell populations at each timepoint. Points overlaid on the histograms indicate the mean mass for each drug condition and timepoint. Horizontal coordinate represents density (dimensions pg^−1^), with the same horizontal scale for all distributions for each model. Some histograms appear narrow because at least one sample for that model accumulated many particles of the same size, compressing the horizontal (density) scale; this is common for TMZ-responsive models (bottom), due to the accumulation of low-mass particles (<25 pg) in the culture as a result of drug-induced cell death.

### Biophysical drug susceptibility testing is consistent with known predictors of TMZ sensitivity

As a first step for evaluating whether biophysical drug susceptibility testing predicts patient response to TMZ, we asked whether the SMR mass assay, the IncuCyte assay, and the CellTiter-Glo assay were consistent with known predictors of TMZ sensitivity. The best-known predictors of TMZ sensitivity are methylation of the O^6^-methylguanine-DNA methyltransferase (MGMT) promoter and mismatch repair (MMR) pathway status *(28)*. Promoter methylation at the MGMT locus leads to reduced RNA and protein expression and an MGMT deficient state, and patients with methylated MGMT promoter are more likely to respond to TMZ and have longer overall survival due to reduced activity of the MGMT DNA repair activity *(29)*. Similarly, unmethylated MGMT promoter is generally associated with a proficient MGMT pathway and treatment induced DNA damage is rapidly repaired in cancer cells, leading to TMZ resistance and shorter patient survival. Therefore, we expected that functional testing would generally detect greater TMZ responsiveness in models established from MGMT-methylated patients, and reduced TMZ responsiveness in models established from MGMT-unmethylated patients. However, in MGMT-deficient tumor cells, MMR deficiency (e.g. via mutation) can also lead to TMZ resistance, as exposed cells can continue to cycle while accumulating a high mutational burden induced by the drug *(28)*. Consistently, the three MMR-deficient models in our cohort showed low functional TMZ responsiveness by all three assays (**Supplemental Fig. 4**). Therefore, to isolate MGMT as a response correlate, we limited our analysis to the subset of 67/70 models which were MMR-proficient (**Fig. 5A**).

**Figure 5.**
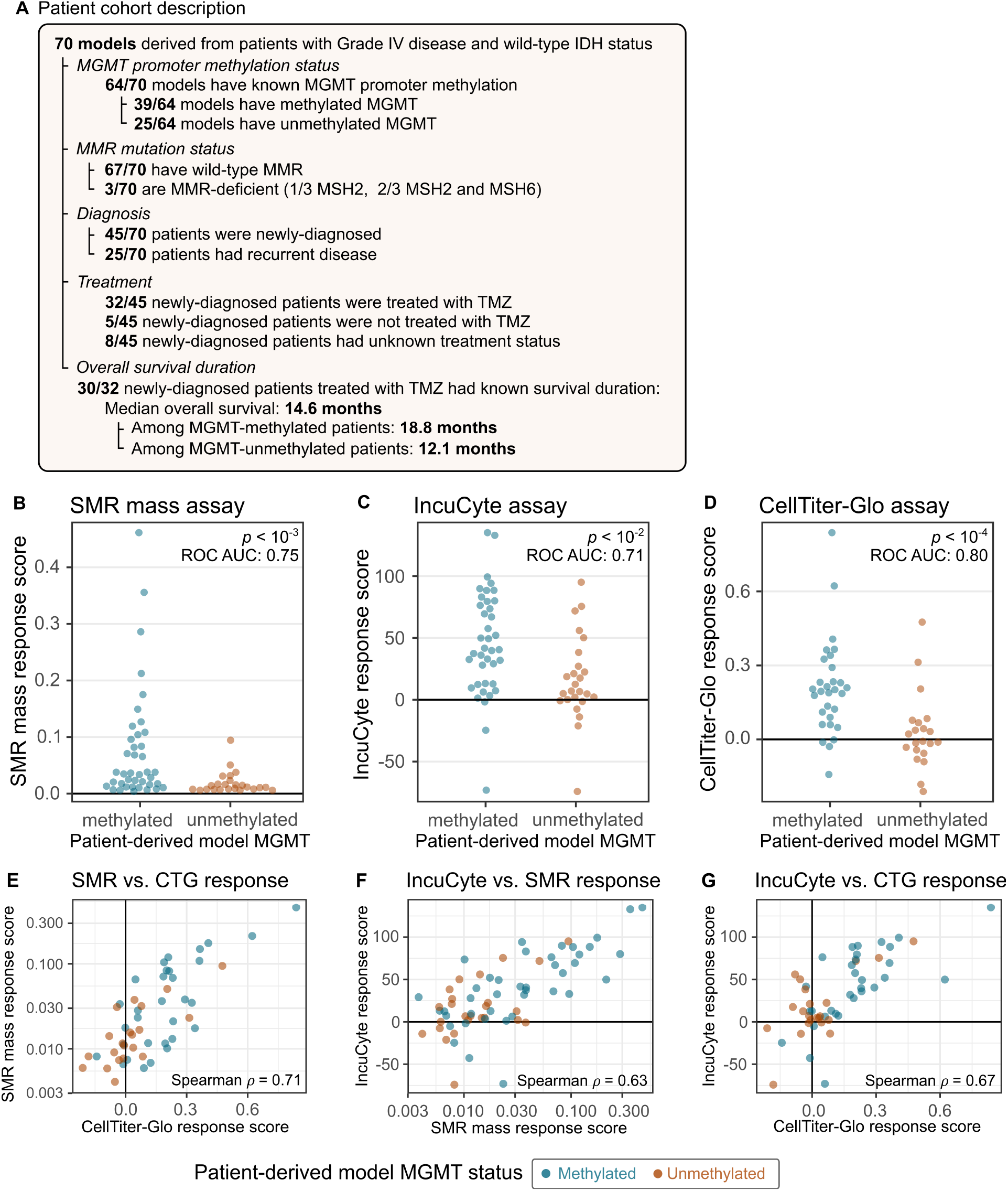
Functional drug susceptibility testing is consistent with MGMT methylation for predicting TMZ sensitivity. **(A)** Clinicopathologic characteristics of the patient cohort from which the patient-derived models were established. **(B-D)** For all three functional assays, MGMT-methylated patient-derived models have significantly higher functional response scores compared to MGMT-unmethylated models: **(B)** SMR mass assay (Wilcoxon rank-sum *p* < 0.001, ROC AUC 0.75), **(C)** IncuCyte assay (Wilcoxon rank-sum *p* < 0.01, ROC AUC 0.71), **(D)** CellTiter-Glo assay (Wilcoxon rank-sum *p* < 0.0001, ROC AUC 0.80). **(E-G)** Correlation between SMR mass response score, CellTiter-Glo response score, and IncuCyte response score across models (shown for models with known MGMT status).

We compared the SMR mass response score, IncuCyte response score, and CellTiter-Glo response score for MGMT methylated versus unmethylated models, and found that for all three functional assays, MGMT-unmethylated models had significantly lower functional TMZ response scores than methylated models (**Fig. 5B-D**). To compare the degree to which each functional biomarker differed between MGMT-methylated and MGMT-unmethylated models, we computed the receiver operator characteristic area-under-the-curve statistic (ROC AUC), interpretable as the probability that a randomly-chosen MGMT-methylated sample will have a larger TMZ response score than a randomly-chosen MGMT-unmethylated sample; the values were comparable for all three functional assays (ROC AUC values: SMR mass assay 0.75, IncuCyte assay 0.71, CellTiter-Glo assay 0.80).

For 59/69 of the patient-derived neurosphere models in our cohort with known MGMT model status (model MGMT; mMGMT), we also independently measured or collected from clinical records the MGMT promoter methylation status of the primary patient tumor sample (patient MGMT; pMGMT). Both mMGMT and pMGMT had similar correlations with functional response scores (**Supplemental Note 2)**. Overall, these results suggest that functional drug sensitivity testing is broadly consistent with the best currently available predictors of TMZ sensitivity.

Next, we asked whether the SMR, IncuCyte, and CTG functional assays measured similar TMZ responsiveness for each patient-derived model. Across all 70 patient-derived models, the SMR mass response score, the IncuCyte response score, and the CellTiter-Glo response score were mutually correlated (**Fig. 5E-G**, shown for the 64/70 models with known MGMT status; Spearman correlations 0.71 for SMR vs. CellTiter-Glo; 0.63 for IncuCyte vs. SMR, and 0.67 for IncuCyte vs. CellTiter-Glo). Despite the high degree of correlation between the three functional assays, a subset of patients registered much higher or lower functional response scores on one assay compared to the others. To determine whether these outliers could be attributed to technical or experimental errors in one of the measurements, we manually identified a subset of models for which the functional biomarkers disagreed and carefully reviewed the raw data (**Supplemental Fig. 5**). Of the manually-flagged models for which two or more functional assays disagreed with one another, none were clearly attributable to experimental artefacts in the raw data.

### Biophysical functional assays predict overall survival in patients

Next, we asked retrospectively whether functional testing could predict the overall survival of GBM patients treated with TMZ. To limit potentially confounding effects, we further limited our analysis to the 32 patient-derived models derived from patients who at the time were newly diagnosed, IDH-wild-type, were subsequently treated with TMZ, and for whom overall survival was available. The median overall survival for this group was 14.6 months.

We used the receiver operator characteristic (ROC) to assess the sensitivity and specificity with which each functional biomarker predicted overall survival of 15 months or greater (**Fig. 6A**). For each functional biomarker, we computed the ROC area-under-the-curve statistic (ROC AUC) as a measure of the biomarker’s predictive power. Both the IncuCyte and SMR biophysical assays were moderately predictive of 15-month survival (ROC AUC statistic 0.78 for both the SMR and IncuCyte biophysical assays), while the CellTiter-Glo assay was somewhat less predictive of 15-month survival (ROC AUC statistic 0.66). Predictive power was similar for other binary survival outcomes, i.e., 12-month, 15-month, 18-month, 21-month, and 24-month survival (**Supplemental Fig. 6**).

**Figure 6.**
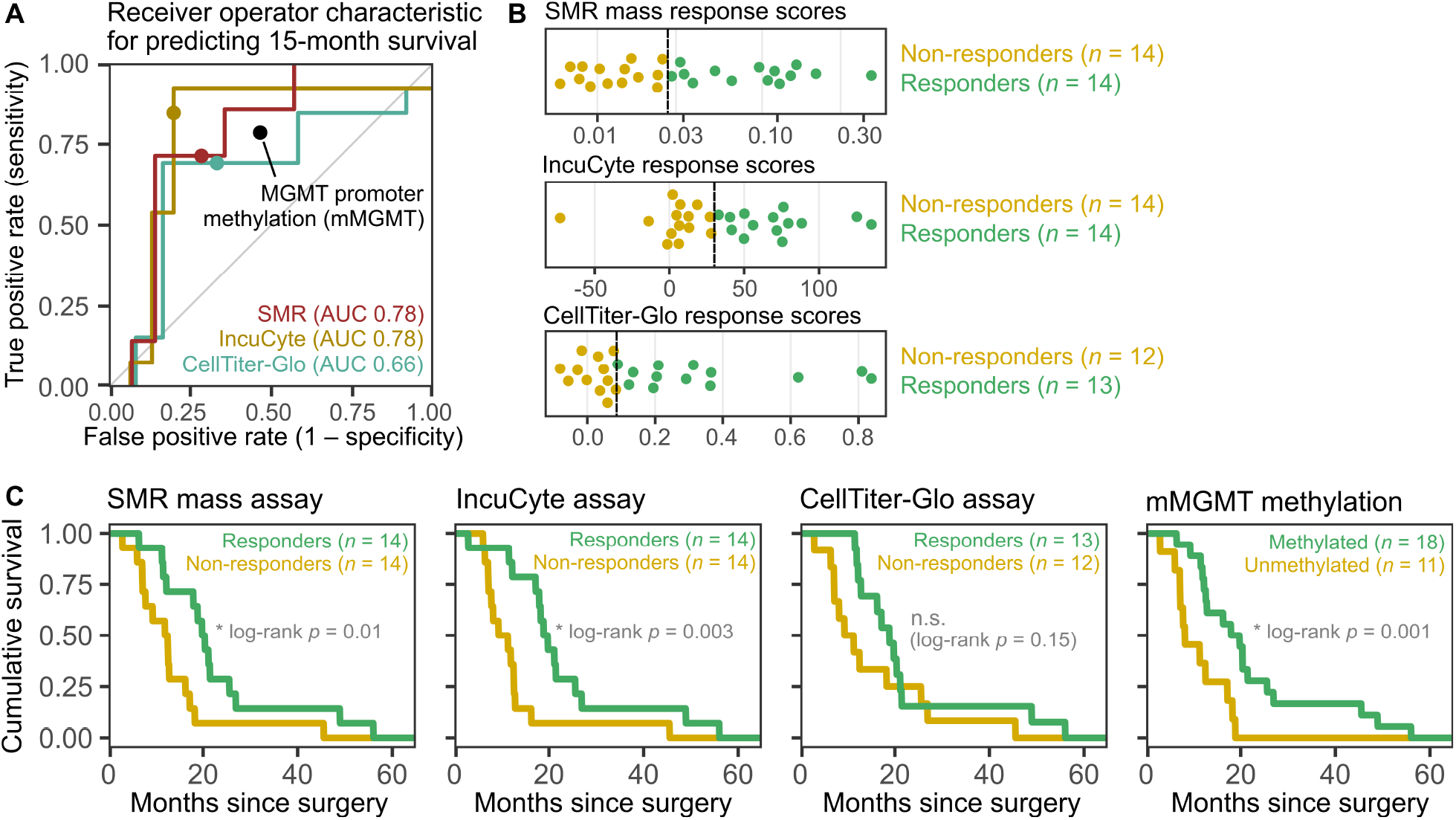
Biophysical functional testing predicts overall survival duration. **(A)** Receiver operator characteristic (ROC) between continuous functional biomarkers (the SMR mass response score, IncuCyte response score, and CellTiter-Glo response score) and a binary survival outcome (15-month survival). Points indicate the performance of the thresholds used for the classification in (B) and (C). (**B)** For each assay, patients within the top 50% of response scores were labeled as functional “responders”, and the bottom 50% were labeled as “non-responders”. **(C)** Overall survival distributions of functional responders versus non-responders. For the SMR and IncuCyte biophysical assays, functional TMZ responders survived significantly longer on therapy than TMZ non-responders (SMR mass assay: median overall survival 20.1 vs. 12.2 months, log-rank *p* = 0.01; IncuCyte assay: median overall survival 19.3 vs. 10.3 months, log-rank *p* = 0.003). Median survival was longer for CellTiter-Glo assay responders than for non-responders (median overall survival 18.8 vs. 12.3 months), but overall survival distributions were not significantly different between these groups (log-rank *p* = 0.15). The predictive power of the SMR mass assay and the IncuCyte assay was comparable to the predictive power of mMGMT promoter methylation (median overall survival 18.8 vs. 8.2 months for mMGMT-methylated vs. MGMT-unmethylated patients; log-rank *p* = 0.001).

To further assess whether functional drug susceptibility testing predicts overall survival, we compared survival distributions between TMZ responsive and non-responsive patients. We chose to classify the 50% of patients with the highest functional response scores as TMZ responders (**Fig. 6B**), since previously-published imaging results suggest that approximately half of glioma patients respond to TMZ *(30, 31)*. While the exact fraction of TMZ-responsive patients is debated, we noted that our results were robust when different fractions of patients were classified as TMZ responders (in the range of 25-75% responders, which generally includes the clinically-estimated range of responses to TMZ; **Supplemental Fig. 7**).

Using this clinical hypothesis-based classification, overall survival was significantly longer for SMR mass assay responders than for SMR mass assay non-responders (**Fig. 6C**; median overall survival 20.1 vs 12.2 months; log-rank *p* = 0.01), and for IncuCyte responders compared to IncuCyte non-responders (**Fig. 6C**; median overall survival 19.3 vs. 10.3 months; log-rank *p* = 0.003). While CellTiter-Glo responders had longer median survival than CellTiter-Glo non-responders (18.8 vs. 12.3 months), there was not a statistically-significant difference in overall survival distributions between these two groups (**Fig. 6C**; log-rank *p* = 0.15). The predictive power of the SMR and IncuCyte biophysical assays was comparable to that of patient-derived model MGMT promoter methylation status (mMGMT; **Fig. 6C**; log-rank *p* = 0.001; median survival 18.8 months vs. 8.2 months for patients with methylated vs. unmethylated mMGMT).

To confirm that the SMR, IncuCyte, and mMGMT biomarkers did not reach statistical significance due only to their increased sample size relative to the CellTiter-Glo (28 eligible patients had SMR and IncuCyte data and 29 patients had known mMGMT status, while only 25 patients had CellTiter-Glo data), we repeated this analysis limited to the subset of 25 patients with CellTiter-Glo data and found similar results. Specifically, among this subset of patients, overall survival was still significantly longer for SMR assay responders versus non-responders (median overall survival 20.4 vs. 12.5 months, log-rank *p* = 0.046), for IncuCyte responders versus IncuCyte non-responders (median overall survival 19.7 vs. 11.3 months, log-rank *p* = 0.01), and for patients with MGMT methylated versus unmethylated models (median overall survival 19.7 vs. 11.3 months, log-rank *p* = 0.01), but not for CellTiter-Glo responders versus non-responders (median overall survival 18.8 vs. 12.3 months, log-rank *p* = 0.15).

Next, we asked whether functional biomarkers could be combined with the MGMT promoter methylation biomarker to make more accurate predictions of patient survival outcomes. In addition to evaluating the SMR, IncuCyte, and mMGMT biomarkers individually (**Supplemental Fig. 8A**), we defined new predictors of TMZ sensitivity based on combinations of these biomarkers. Specifically, we computed the survival distributions of patients who were (1) SMR responders *and* mMGMT-methylated; (2) SMR responders *or* mMGMT-methylated; (3) IncuCyte responders *and* mMGMT-methylated; and (4) IncuCyte responders *or* mMGMT-methylated (**Supplemental Fig. 8B-D**). As expected, the biomarkers requiring both a functional response *and* methylated mMGMT to predict TMZ sensitivity were more specific for identifying TMZ-responsive patients, with patients meeting both criteria having longer overall survival (**Supplemental Fig. 8B**). Similarly, the biomarkers requiring only a functional response *or* methylated mMGMT were more sensitive for identifying TMZ-responsive patients (**Supplemental Fig. 8C**). In practice, the desired balance between sensitivity and specificity for identifying TMZ sensitivity versus resistance will depend on the specific clinical application.

## Discussion

This retrospective study demonstrates that biophysical assays can be used as a functional readout for *ex vivo* drug susceptibility testing, and that in particular, the SMR and IncuCyte biophysical assays can be used in individual patients to predict overall survival duration following TMZ treatment, with predictive power comparable to MGMT promoter methylation.

One interesting finding from our work was that all three functional assays identified a continuous spectrum of TMZ responsiveness in the patient-derived neurosphere models, rather than a bimodal distribution with clearly distinct TMZ-responsive and TMZ-resistant models. This result suggests that GBM patients may also exhibit a continuous spectrum of cell-intrinsic TMZ sensitivity versus resistance, rather than the binary classification typically represented by the presence or absence of MGMT promoter methylation. The notion of a continuous spectrum of TMZ responsiveness is consistent with previous work showing that MGMT promoter methylation status is also continuously distributed across patients, and that there is not a clear, unambiguous level of methylation separating “methylated” from “unmethylated” patients *(32)*. Furthermore, recent studies suggest that quantitative levels of MGMT methylation in gliomas may also correlate with the response to TMZ *(33)*. It is possible that a continuous distribution of functional response scores could be used to make more quantitative predictions of survival for individual patients, by classifying patients into multiple levels of TMZ sensitivity depending on the extent of their functional TMZ response. Assessing whether these approaches are predictive will require larger patient cohorts and greater statistical power. However, if validated, more quantitative, graded predictions of drug susceptibility could be a valuable addition to existing clinical workflows.

As new functional precision medicine approaches move toward clinical implementation, practicality is a key consideration. One key metric is the throughput of the assay. In this study, it took four months to perform functional testing on 70 patient-derived neurosphere models. The main rate-limiting factor in our study was the time required to culture the patient-derived neurosphere models. However, the next rate-limiting factor would have been the throughput of profiling the cultures using the SMR mass assay. The on-device data collection time was approximately 20 minutes per SMR sample; therefore, measuring a total of 12 conditions per patient model (2 drug conditions and 6 timepoints), required a total of 3 hours of SMR instrument time per patient model. This level of throughput is likely compatible with current clinical pathology workflows. Further, we estimate that the time-per-sample could be significantly reduced: as we have shown previously *(20)*, using more concentrated cell samples, the SMR can achieve mass measurement throughput reaching thousands of cells per minute, potentially reducing the required SMR instrument time per sample by several fold.

A second key practical consideration is sample consumption, i.e., the number of tumor cells required to perform functional testing. Depending on the cancer, a typical solid tumor biopsy or resection might contain several hundred thousand tumor cells, setting an upper bound on the number of drug conditions or replicates on which testing can be performed. In our study, because the patient-derived neurosphere models could be propagated *ex vivo* to obtain more sample material, tumor cells were abundant and so we did not fully optimize the study design to use cells efficiently. Despite this, our assays consumed only ~120,000 cells per patient for each drug condition tested, of which ~90,000 were used by the SMR mass assay. These sample requirements are comparable to next-generation sequencing, which is already a standard component of many clinical workflows and requires 10 ng-1 μg of DNA from typically 10^3^-10^5^ tumor cells *(34)*. However, the sample input requirements of the SMR mass assay could be reduced by several fold since sufficient statistical power can be obtained by weighing as few as ~2,000 cells per sample, which is considerably less than the 15,000 cells per sample used in the current protocol (**Supplemental Note 1**).

A frequent concern with functional drug susceptibility testing approaches is that *ex vivo* testing only measures tumor cell-intrinsic drug susceptibility, and does not capture all of the other factors that determine a patient’s overall survival duration on therapy. In particular, our study did not assess the effects of radiation therapy, which most glioblastoma patients receive concurrently with TMZ *(35)*. Depending on the drug and disease, it is possible that synergistic effects could exist between chemotherapy and radiation, and that these effects would not be captured in functional susceptibility testing of the drug alone. Regardless, assessing tumor cell-intrinsic drug susceptibility is still useful, because for most drugs there still exists clear variation in drug susceptibility across patients, including glioma patients treated with TMZ *(31)*. Further, most molecular biomarkers (such as EGFR mutation status in lung cancer, and many other examples) are also used as indicators of tumor cell-intrinsic drug susceptibility, so this limitation is not unique to functional testing.

Although most functional testing approaches have focused on drugs with cell-autonomous mechanisms, similar approaches could likely be extended to other drugs involving tumor-microenvironment interactions by using different *ex vivo* model systems in which tumor cells are co-cultured with other relevant cell types *(3)*.

We found that both the IncuCyte and SMR biophysical assays were consistent with the MGMT biomarker and that both functional assays predicted statistically-significant differences in overall survival between responders and non-responders. It is interesting to note that while the CellTiter-Glo assay was also consistent with the MGMT biomarker, in our hands, there was not a statistically-significant different in overall survival distributions between CellTiter-Glo responders versus non-responders. However, this one data point should not be viewed as an indictment of the CellTiter-Glo assay as a whole, and should be viewed in light of the large body of existing work in which the assay has successfully been used for drug susceptibility testing. As others have described previously *(36)*, even simple experiments such as measuring the response of cells to *in vitro* drug exposure can be affected by factors such as cell counting protocols, micro-titer plate selection, and cell seeding density, all of which result in significant inter-center variability. We ran the CellTiter-Glo assay using one particular assay format (i.e., plate type, cell seeding density, feeding schedule, drug dose, and timepoint selection), and while our selection was a reasonable choice, it is entirely possible that under different conditions the assay could be more predictive. Further studies might reveal biological underpinnings that explain the differences in drug responsiveness using biophysical versus metabolic assay readouts.

In addition to our focus on using biophysical measurements for drug susceptibility testing, a key distinction between this work and most other applications of functional drug susceptibility testing is that this study focuses on stratifying patients by predicted susceptibility to one specific drug, as opposed to selecting between several candidate drugs. The most common application of functional drug susceptibility testing is: given a patient and several candidate drugs, which drug is likely to be most effective? Previous studies in leukemia *(37)* and multiple myeloma *(6)* have focused on this question, where the goal is to test many candidate drugs against a patient’s tumor, and select the most effective from among them. However, our study focused on a different question: given a single standard-of-care drug and a population of patients, which patients are most likely to benefit from the drug? While more limited in scope compared to studies that test dozens of novel drugs and combinations, such studies focused on single clinical scenarios of need are potentially more readily translated into clinical adoption.

## Materials and Methods

### Description of patient cohort

All patients, clinical data and models were studied following consent to research (DFCI IRB#10-417) or waiver of consent (DFCI IRB#10-043) per institutional review board procedures at the Dana-Farber Cancer Institute and Brigham and Women’s Hospital. We performed functional testing on 70 patient-derived models established from GBM patients, all of whom had wild-type IDH. Of these 70 models, functional response data was successfully collected for 68/70 models using the SMR assay, 55/70 models using the CellTiter-Glo assay, and 68/70 models using the IncuCyte assay. Of the 70 models, 67/70 have a wild-type mismatch repair (MMR) phenotype, while 3/70 have MMR mutations (1/3 MSH2 mutation, 2/3 both MSH2 and MSH6 mutations).

For comparing functional response data to MGMT methylation status, we restricted our analysis to the subset of patients with wild-type MMR. Of these 65 patients, 64/65 had models with known MGMT methylation status (39/64 methylated, 25/64 unmethylated). Further, of these 65 patients, 63/65 had SMR mass assay data, 50/65 had CellTiter-Glo assay data, and 63/65 had IncuCyte assay data.

For comparing functional response data to patient survival outcomes, we further restricted our analysis to patients who were newly-diagnosed and were treated with TMZ. Of the 70 models, 45/70 were newly-diagnosed, and of these, 43/45 had wild-type MMR. Of the 43 models established from newly-diagnosed patients with wild-type MMR, 32/43 were treated with TMZ, 4/43 were not treated with TMZ, and the remaining 7/43 had unknown TMZ treatment status.

Of the 32 eligible models (those established from newly-diagnosed patients with wild-type MMR who were treated with TMZ), overall survival duration was known for 30/32 patients. Of these 30 patients, 28/30 have SMR mass assay data, 28/30 have IncuCyte assay data, 25/30 have CellTiter-Glo assay data, and 29/30 have known patient-derived model MGMT status (18/29 methylated, 11/29 unmethylated).

### GBM patient-derived model initiation and culture

GBM patient-derived models were established and maintained as described previously *(19)*. Briefly, GBM tumor resections were subjected to enzymatic and mechanical dissociation, then seeded in neurosphere culture conditions. The models were propagated in a proprietary neurosphere culture medium (NeuroCult; STEMCELL Technologies) supplemented with additional growth factors (20 ng/mL epidermal growth factor, 10 ng/mL fibroblast growth factor) and 2 μg/mL heparin. Models were propagated in ultra-low attachment flasks (Corning 3814), and passaged by dissociating to single cells (5-minute treatment with 1X Accutase at 37 °C; STEMCELL Technologies) and resuspending at a concentration of 100k-300k cells/mL.

### Functional testing of GBM patient-derived models

For the SMR assay, patient-derived models were dissociated to single cells (5-minute treatment with 1X Accutase at 37 °C), then seeded at a density of 15,000 cells/mL in 24-well ultra-low attachment plates (Costar 3473), with a volume of 1 mL/well. After allowing 24 hours for neurosphere formation, the cells were exposed to 20 μM temozolomide or a vehicle control (0.1% DMSO). Medium was replenished (100 μL/well) at days 3, 6, 10, and 13 after drug exposure, while cells were sampled for SMR measurement on days 3, 5, 7, 10, 12, and 14. For SMR mass measurement, neurospheres were dissociated to single cells by treatment with 1X Accutase for 10 minutes at 37 °C, then resuspended in 50 uL medium and sampled by the SMR for 20 minutes. Samples with fewer than 300 cells detected in this time period were excluded.

For the CellTiter-Glo assay, patient-derived models were dissociated to single cells (5-minute treatment with 1X Accutase at 37 °C), then seeded at a density of 1500 cells/mL in 96-well ultra-low attachment flat-bottom plates (Corning 7007), with a volume of 100 μL/well, with three biological replicates per condition. After allowing 24 hours for neurosphere formation, the cells were exposed to 20 μM temozolomide or a vehicle control (0.1% DMSO). Medium was replenished (10 μL/well) at days 3, 6, 10, and 13 after drug exposure. At days 3, 5, 7, 10, 12, and 14 after drug exposure, the CellTiter-Glo assay was performed following the manufacturer’s protocol. For the CellTiter-Glo assay, we excluded timepoints for which there was unusually high variation between replicates (specifically, we excluded timepoints for which the coefficient of variation of the measured luminescence signal between replicates was greater than 30%, excluding a total of 7% of the measured timepoints).

For the IncuCyte assay, patient-derived models were dissociated to single cells (5-minute treatment with 1X Accutase at 37 °C), then seeded at a density of 1500 cells/100 μL in 96-well ultra-low attachment round-bottom plates (CLS7007), with three biological replicates per condition. After allowing 24 hours for neurosphere formation, the cells were exposed to 20 μM temozolomide or a vehicle control (0.1% DMSO). Cells were then loaded on an IncuCyte S3 Live-Cell Analysis Imaging System (Essen BioScience), and imaged every 2 hours for 14 days following drug exposure, using the built-in Spheroid Analysis module. Images were segmented to identify neurospheres, then spheroid size was quantified as the total projected spheroid area per well. To eliminate cases in which the IncuCyte did not successfully capture an in-focus image of the neurosphere, we excluded frames in which the total observed spheroid area per well was less than 0.25 mm^2^. For the IncuCyte assay, replicates were excluded from analysis by reviewing the images and excluding wells for which (1) the spheroid was not visible in the image, e.g., due to an auto-focus failure with the automated microscope, or (2) the built-in image segmentation algorithm did not accurately identify the boundaries of the spheroid.

### Functional testing of Ba/F3 BCR-ABL and Ba/F3 BCR-ABL T315I cell lines

Ba/F3 BCR-ABL and Ba/F3 BCR-ABL T315I cell lines were a gift from the Weinstock laboratory at the Dana-Farber Cancer Institute. Both the Ba/F3 BCR-ABL and Ba/F3 BCR-ABL T315I cell lines were maintained in RPMI-1640 medium (ThermoFisher) supplemented with 10% FBS (Sigma-Aldrich), 25 mM HEPES (Gibco), and 1X antibiotic/antimycotic (Gibco). For maintenance, cells were passaged every 2-3 days to a minimum concentration of 75,000 cells/mL.

For functional testing, cells were seeded at 75,000 cells/mL in 12-well plates (Argos P1012), with a volume of 1 mL/well. After 24 hours, cells were dosed with imatinib (Santa Cruz Biotechnology), ponatinib (Santa Cruz Biotechnology), or a vehicle control (0.2% DMSO). At 8-10 hours of drug exposure, cells were collected and immediately sampled by a SMR system for up to 20 minutes per sample.

### Functional testing of PC9 and PC9-GR4 cell lines

PC9 and PC9-GR4 cell lines were a gift from the Jänne laboratory at the Dana Farber Cancer Institute. Both cell lines were maintained in RPMI-1640 medium (ThermoFisher) supplemented with 10% FBS (Sigma-Aldrich), 25 mM HEPES (Gibco), and 1X antibiotic/antimycotic (Gibco) in 75 cm^2^ culture flasks (VWR 10062-860). For maintenance, cells were passaged every 3 days to a concentration 50k cells/mL in a volume of 15 mL (i.e., 10k cells/cm^2^). At each passage, medium was aspirated, cells were detached from the culture surface by incubating with 0.25% trypsin-EDTA (ThermoFisher) for 10 minutes at 37 °C, then washing with medium and resuspending at the desired concentration.

For functional testing, cells were trypsinized and seeded in 12-well plates (Argos P1012) at a concentration of 50k cells/mL in a volume of 1 mL. After 24 hours, cells were exposed to gefitinib (Selleck Chemicals), osimertinib (Selleck Chemicals), or a vehicle control (0.1% DMSO). At timepoints of 8, 16, and 24 hours of drug exposure, cells were trypsinized then resuspended in 150 μL medium for SMR measurement, with a measurement duration of 20 minutes per sample.

### SMR mass response score

For each functional assay, a “response score” was calculated that summarizes the extent to which the sample responds to the drug treatment compared to a vehicle control across all measured timepoints.

The SMR mass response score is based on the Hellinger distance between the control and drug-treated cell mass distributions. The Hellinger distance is a measure of statistical distance between mass distributions *P*(*m*) and *Q*(*m*), and is defined as:

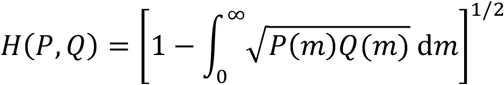

A Hellinger distance of 0 corresponds to no difference between the control and drug-treated mass distributions (i.e., no drug response), and a Hellinger distance of 1 would correspond to completely non-overlapping mass distributions. This summary statistic has the advantage that is agnostic to whether drug treatment caused mass to increase or decrease, and simply identifies the degree of change in the mass distribution. We evaluated the Hellinger distance between each control-drug pair by computing kernel density estimates of *P* and *Q* and then numerically integrating. Further, we obtained bootstrap standard errors and confidence intervals for the Hellinger distance by repeatedly resampling the measured cell mass distributions and re-computing the kernel density estimates and Hellinger distance (200 iterations per sample).

For the GBM patient-derived models, the SMR mass response score is defined as the average of the Hellinger distance between the TMZ and vehicle control samples at days 5, 7, and 10.

### CellTiter-Glo response score

For the GBM patient-derived models, the “CellTiter-Glo response score” was defined as 1 − (average viability across timepoints), where “viability” is defined as the ratio of the mean CTG luminescence signal for the TMZ sample to the mean CTG luminescence signal for the vehicle control sample. This metric integrates the difference between the treatment and control conditions across all measured timepoints, with larger CTG response scores corresponding to a greater reduction in CTG luminescence signal in the drug-treated samples compared to matched controls.

### IncuCyte response score

For the GBM patient-derived models, IncuCyte images were segmented to estimate the total spheroid area per well. We excluded images in which the measured spheroid area was less than 0.25 mm^2^, which typically correspond to cases in which the spheroid was out of focus. We filtered the spheroid size traces using locally-estimated scatterplot smoothing (LOESS). Finally, the “IncuCyte response score” was defined by integrating the difference in spheroid size between the vehicle control and drug-treated conditions, averaging across replicates, and normalizing to the growth rate of untreated cells. Larger IncuCyte response scores correspond to a greater reduction in size in the drug-exposed samples compared to matched controls.

### SMR operation

SMR mass measurements were performed using parallel SMR arrays, as described previously *(20)*. Briefly, single cells in suspension flow through a micromechanical resonator with an embedded fluidic channel, generating a shift in resonance frequency proportional to the cell’s mass. The sensors are calibrated by measuring monodisperse polystyrene beads of known mass. The measurements described here were performed using a parallel SMR array, in which twelve sensors are connected fluidically in parallel and operated simultaneously for increased throughput *(20)*. Before measurement, the fluidic channels were passivated with poly-L-lysine-grafted poly(ethylene glycol). Cell samples remained at room temperature throughout the (typical) 20-minute measurement duration.

### Receiver operator characteristic analysis

To evaluate whether functional biomarkers predicted patient outcome, we computed the receiver operator characteristic (ROC) between the functional biomarker value (i.e., the SMR mass response score, IncuCyte response score, or CellTiter-Glo response score) and the binary survival outcome of interest (e.g., 15-month survival). We computed the ROC area-under-the-curve statistic (ROC AUC) by integrating the receiver operator characteristic true positive rate with respect to false positive rate.

### Measurement of MGMT promoter methylation

Methylation of the CpG island of the MGMT gene was measured using standard methylation-specific PCR at the Brigham and Women’s Hospital Center for Advanced Molecular Diagnostics. Specifically, bisulfite treatment converted unmethylated (but not methylated) cytosines to uracil, prior to PCR using primers specific for either the methylated or modified unmethylated DNA *(38)*. PCR products were analyzed using capillary gel electrophoresis in duplicate parallel runs. Partially methylated or ambiguous calls were verified by a molecular pathologist (A.K.) and binned into one of the categories based on this review.

## Acknowledgments

The authors thank David Weinstock for providing the Ba/F3 BCR-ABL T315I cell line, and Pasi Jänne for providing the PC9 and PC9-GR4 cell lines.

## Author contributions

M.A.S., S.M, S.R.M., K.L.L designed the experiments. M.A.S., S.M., J.Y., M.M. performed the experiments. S.M., P.W., A.K. and J.G. collected and interpreted patient clinical and molecular data. S.M., M.T., and K.C. established patient-derived neurosphere models. M.A.S. and S.M. analyzed the data. M.A.S. wrote the manuscript with contributions from S.M., K.L.L., and S.R.M. All authors reviewed and approved the manuscript. M.A.S. and S.M. contributed equally to this work. K.L.L. and S.R.M. share senior authorship.

## Supplemental Note 1: Comparing throughput of the MAR and SMR mass assays

For both the MAR assay and the SMR mass assay, throughput (in samples measured per hour of instrument time) is determined from the rate at which individual cells can be measured and the number of cells required to detect a significant MAR or cell mass changes with sufficient statistical power.

Previously, it has been reported that serial SMR array devices measure MAR at a maximum rate of approximately 200 cells/hr *(39)*. While required sample sizes depend strongly on the specific cell type and the expected effect size of the drug, previous studies using MAR have typically reported sample sizes on the order of 100-200 cells *(6, 19, 39)*. Therefore, we estimated that the MAR assay has average throughput of 1 sample per hour of instrument time.

We used a similar approach to estimate the sample throughput of the parallel SMR array. Based on a previous study, these devices can measure cell mass at a maximum throughput of 6800 particles/minute when optimized for speed *(20)*. For our order-of-magnitude throughput calculations we used a more conservative estimate of 1000 particles/minute, since most tumor cell samples are sparse and throughput is lower when measuring these generally lower-concentration samples. Next, to estimate required sample sizes, we computed the number of measurements required to detect a significant reduction in mean mass with specified power (1 − *β*), and confidence (1 − *α*), obtaining the following expression for a drug treatment that reduces mean cell mass from *m* to *m* − Δ*m*:

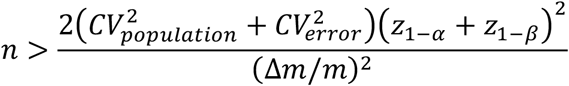

where *n* is the required sample size, *CV*_*populatin*_ is the coefficient of variation of the untreated cells’ masses, and *CV*_*error*_ is the fractional error of the mass measurement, and *z* is the inverse normal cumulative distribution function. Although we generally compared drug-treated versus control samples using the Hellinger distance rather than directly comparing the mean mass, this calculation provides a rough estimate of required sample sizes. Using this expression, a sample size of *n* = 2000 cells allows us to detect a reduction in mean cell mass as small as 3% for a typical sample (*CV*_*populatin*_ = 30%, *CV*_*error*_ = 1%) with 80% power and 99% confidence. Therefore, we estimated that the parallel SMR array has average throughput of 1 sample per 2 minutes of instrument time.

## Supplemental Note 2: Comparing patient-derived model MGMT status to patient MGMT status

We measured the MGMT methylation status of the patient-derived neurosphere models (“mMGMT”) for 69/70 models in our cohort. For 59/69 of these patients, we also independently measured MGMT methylation status of the primary tumor sample at the time of collection (“pMGMT”). Here we compare in detail for which patients these two MGMT status results (mMGMT and pMGMT) agree and disagree.

Of the 59 models where both pMGMT and mMGMT are known, 35/59 (59%) of models had methylated mMGMT and the remaining 24/59 (41%) had unmethylated mMGMT. Of the 35 models with methylated mMGMT, 19/35 (54%) also had methylated pMGMT, while 16/35 (46%) had unmethylated pMGMT. Of the 24 models with unmethylated mMGMT, 17/24 (71%) also had unmethylated pMGMT, while 7/24 (29%) had methylated pMGMT.

In other words, 19/59 patients (32%) had both methylated pMGMT and methylated mMGMT; 17/59 patients (29%) had both unmethylated pMGMT and unmethylated mMGMT; 7/59 patients (12%) had methylated pMGMT but unmethylated mMGMT; and 16/59 patients (27%) had unmethylated p MGMT but methylated mMGMT.

It remains unclear whether these shifts in MGMT methylation between the patient sample and patient-derived model occur due to bias in the MGMT assay (either an abundance of false negative unmethylated calls from primary samples, or false positive methylation calls from PDM samples), or whether this corresponds to a real phenotypic shift between the primary sample and patient-derived model.

While the pMGMT and mMGMT biomarkers were both correlated with functional testing results (i.e., MGMT methylated samples tended to have more significant TMZ responses), the pMGMT biomarker was slightly more strongly correlated than was the mMGMT biomarker. We computed the ROC AUC statistic to quantify to what extent functional response scores are segregated by both the pMGMT and mMGMT biomarkers. For all three assays, the ROC AUC statistic was similar for pMGMT and mMGMT (IncuCyte, ROC AUC 0.67 vs. 0.71; SMR, ROC AUC 0.79 vs. 0.75; CellTiter-Glo, ROC AUC 0.82 vs. 0.80).

**Supplemental Figure 1.**
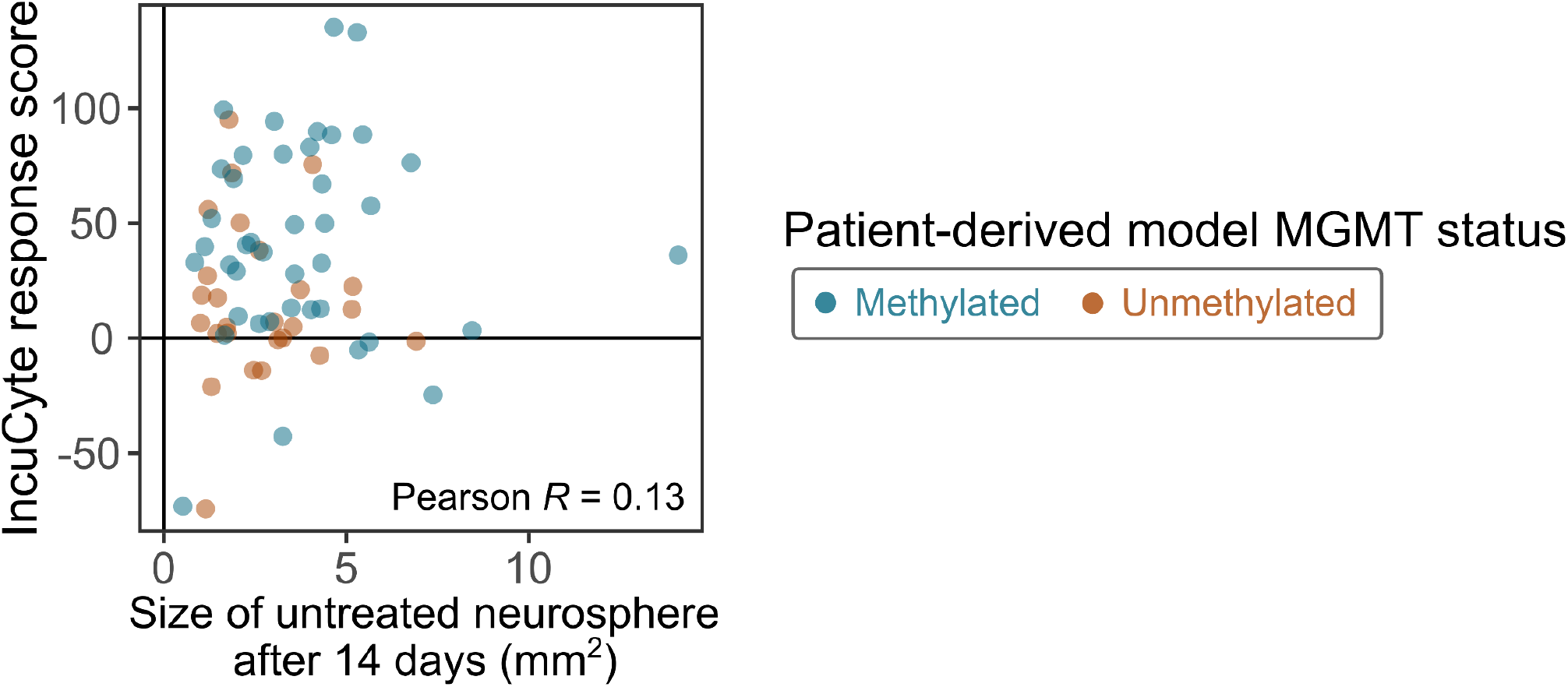
The IncuCyte response score is uncorrelated with the growth rate of untreated neurospheres.

**Supplemental Figure 2.**
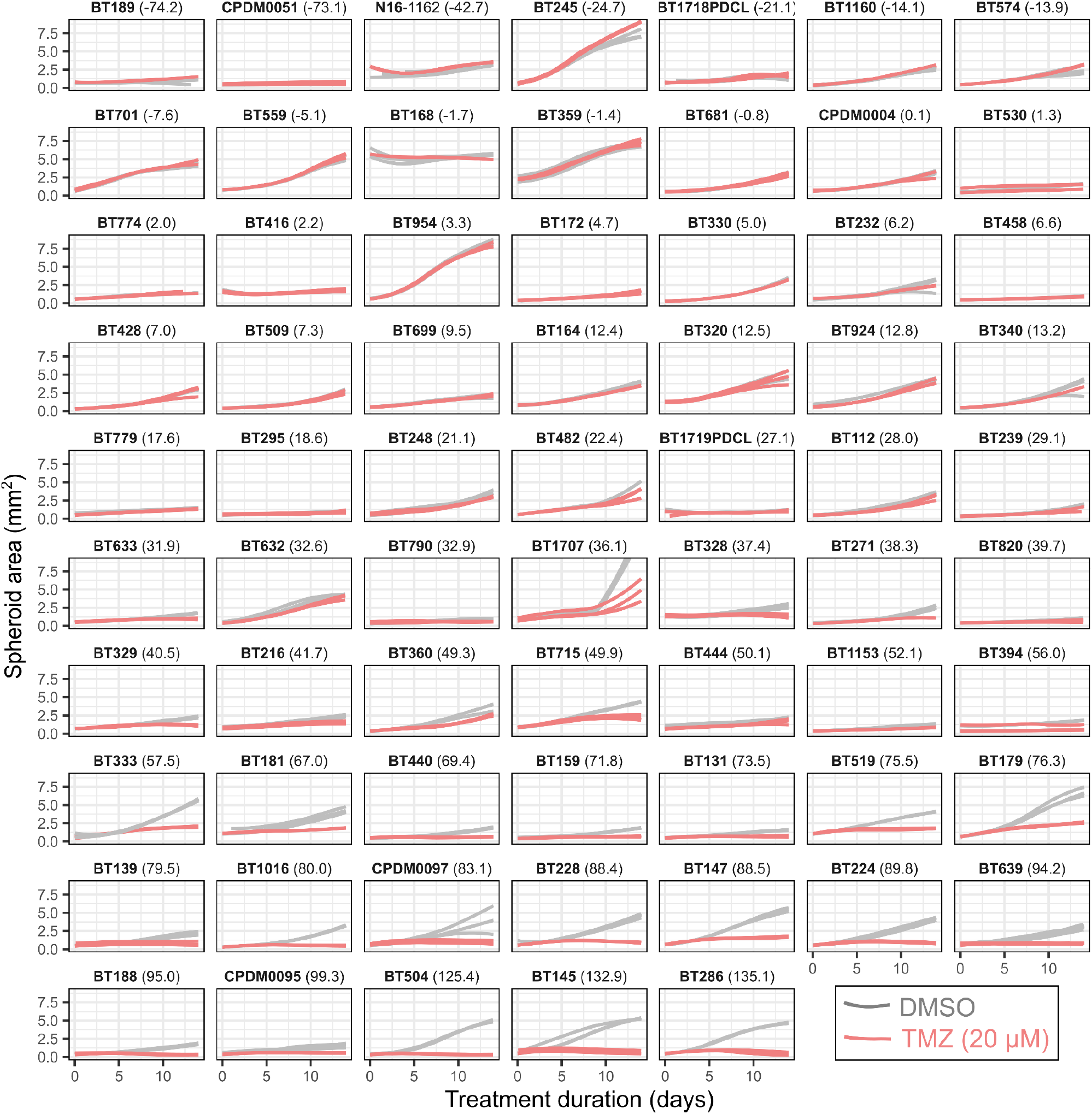
Functional drug susceptibility testing using the IncuCyte assay, which tracks spheroid growth based on microscopy. The 68 patient-derived neurosphere models are arranged in order of low to high IncuCyte response score.

**Supplemental Figure 3.**
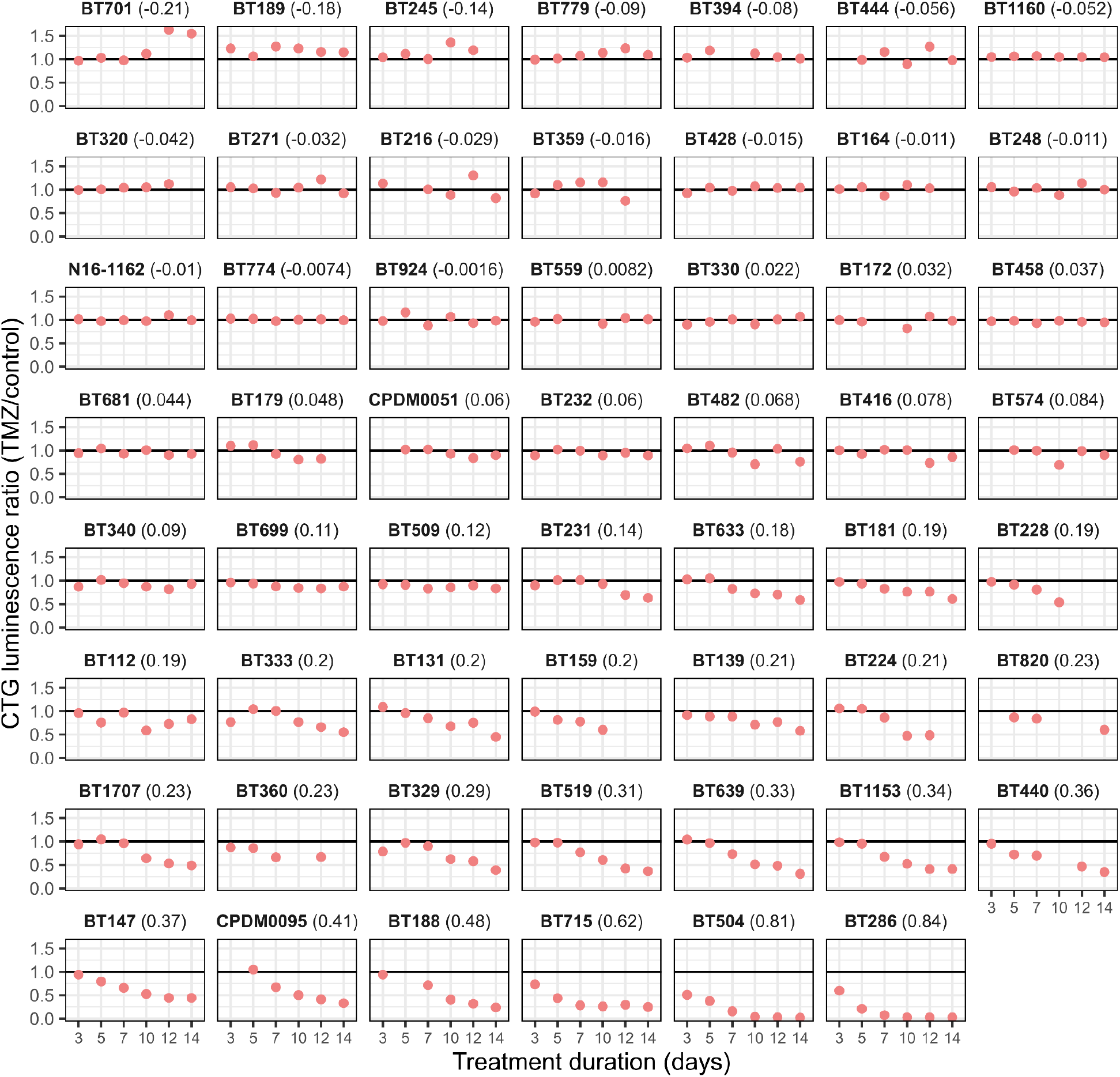
Functional drug susceptibility testing using the CellTiter-Glo assay, which measures ATP levels as a proxy for numbers of viable, metabolically-active cells. The 55 models are arranged in order of low to high CellTiter-Glo response score. We excluded timepoints for which there was unusually high variation between replicates (those for which the coefficient of variation of the measured luminescence signal was greater than 30%, excluding 6% of timepoints with the highest between-replicate variation).

**Supplemental Figure 4.**
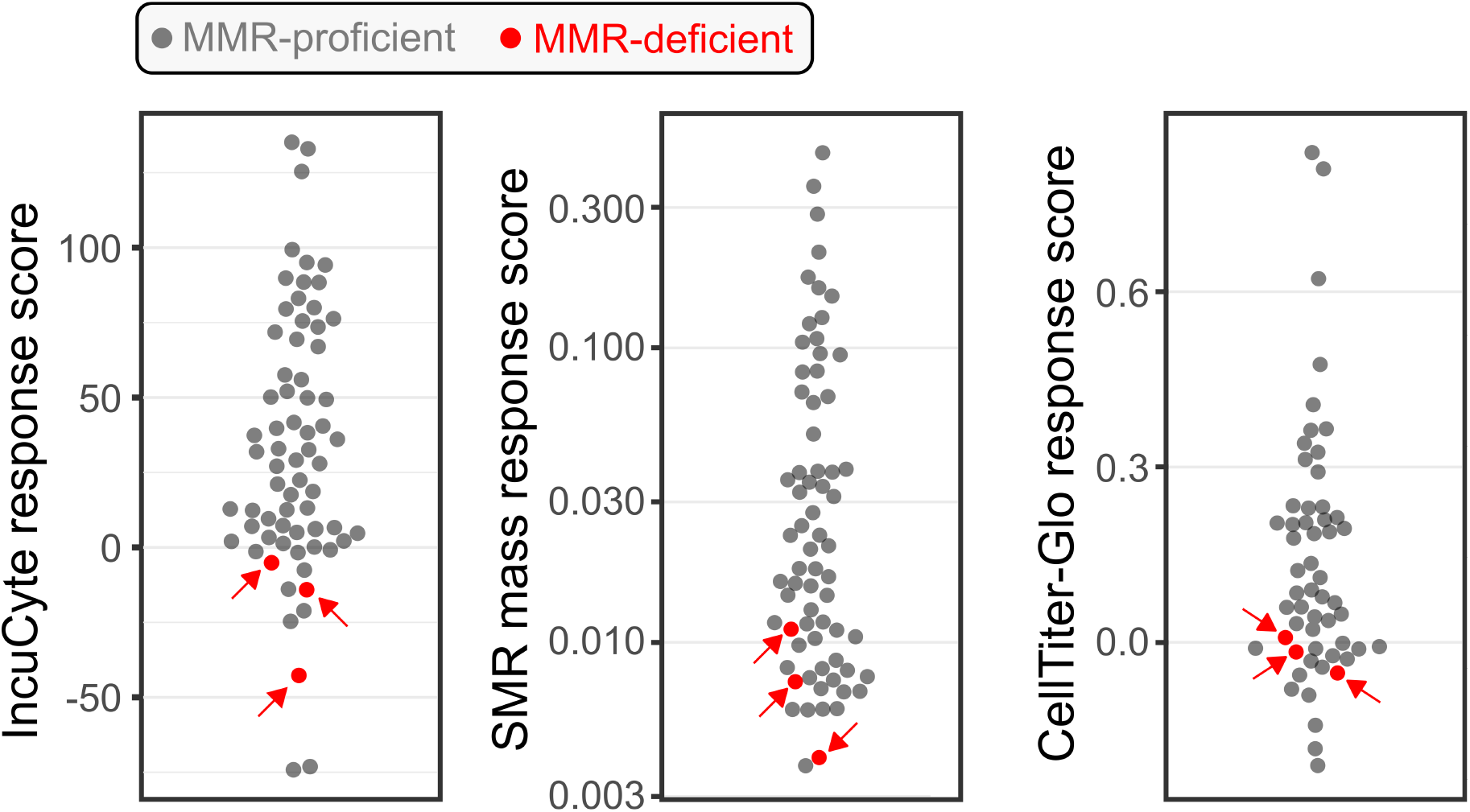
Mismatch-repair-deficient patient-derived neurosphere models are functionally non-responsive to TMZ by all three functional assays.

**Supplemental Figure 5.**
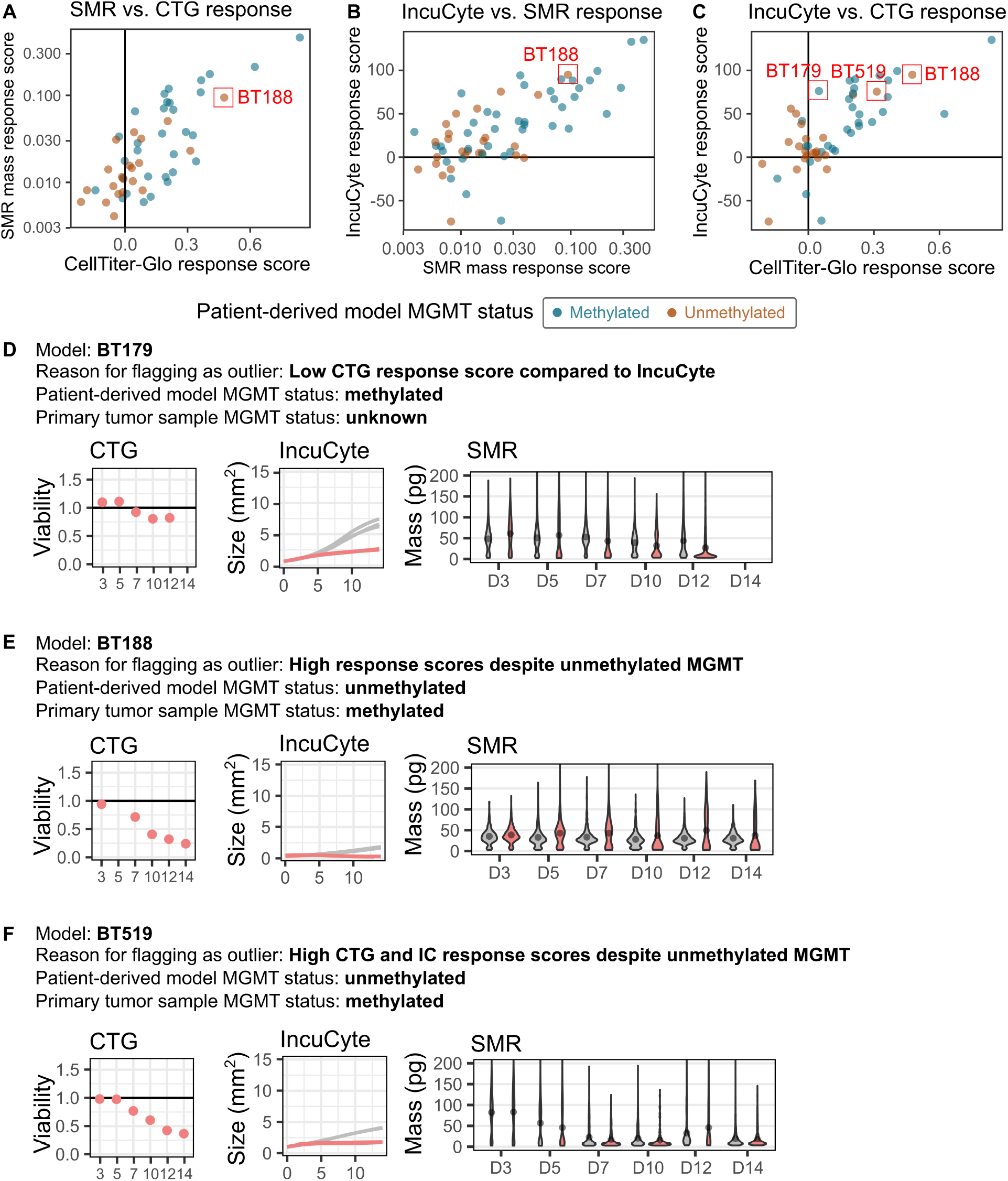
Selected examples of models with differences in predicted TMZ sensitivity across assays. **(A-C)** Functional response scores for the SMR mass assay, IncuCyte assay, and CellTiter-Glo assay. Three patient-derived models (BT179, BT188, BT519) were manually flagged as “outliers” due to different patterns of TMZ responsiveness across assays. **(D)** CTG, IncuCyte, and SMR mass response scores for BT179. BT179 is a strong TMZ responder as measured by the SMR and IncuCyte assays, but a weaker responder as measured by the CellTiter-Glo assay. There were no obvious technical problems with any assay that could explain this difference. **(E)** BT188 appeared to be highly TMZ-responsive despite having unmethylated patient-derived model MGMT status. However, this model was one of the 7/24 unmethylated models for which the primary patient sample had methylated MGMT. **(F)** BT519 appeared to be highly TMZ-responsive despite having unmethylated patient-derived model MGMT status. However, this model was one of the 7/24 unmethylated models for which the primary patient sample had methylated MGMT.

**Supplemental Figure 6.**
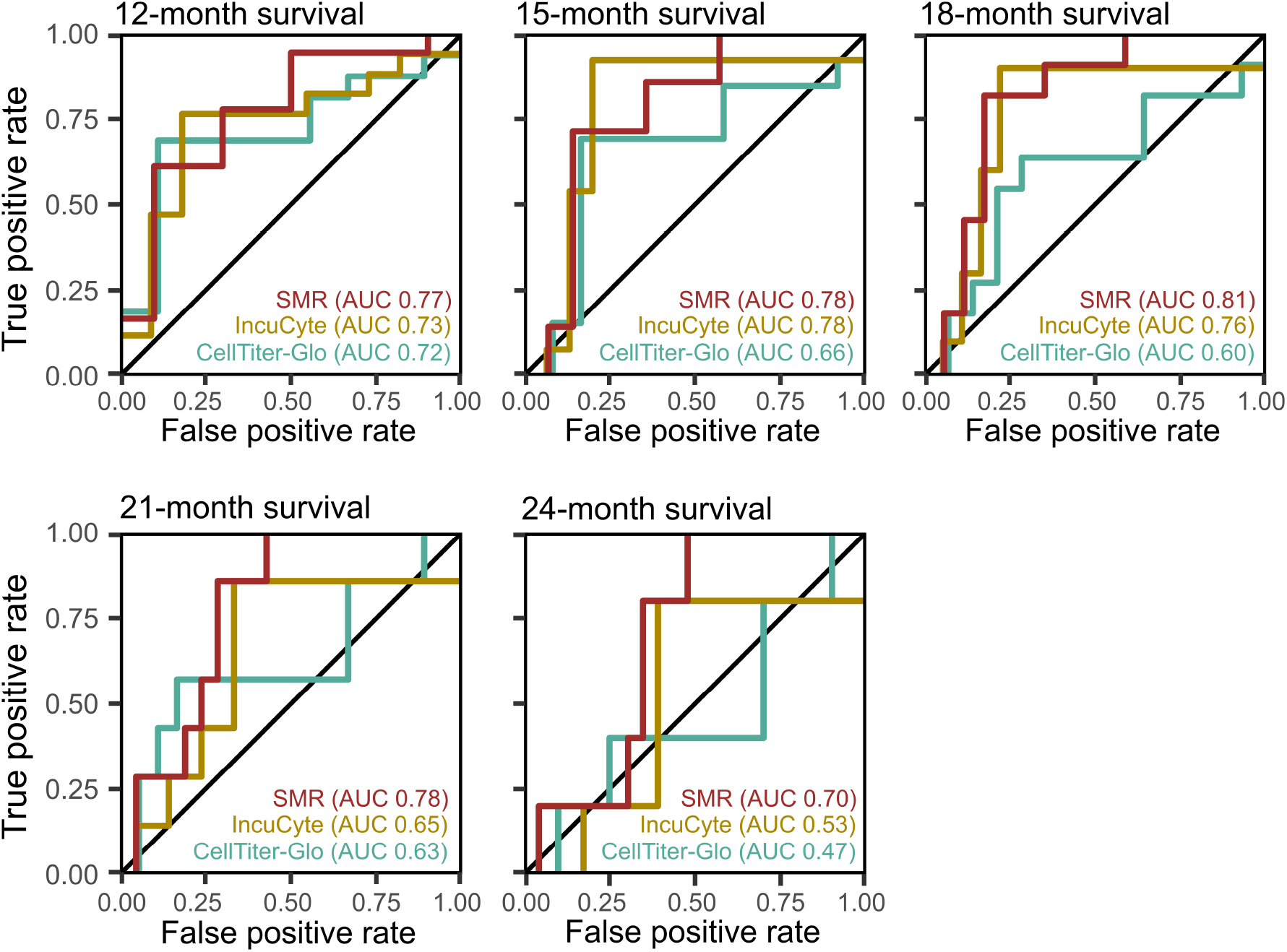
Predicting binary survival outcomes using functional biomarkers. Receiver operator characteristic for predicting 12-, 15-, 18-, 21-, and 24-month survival using the SMR mass assay, IncuCyte assay, and CellTiter-Glo assay. Predictive power (here, quantified as the ROC AUC statistic) is similar regardless of the binary survival outcome used to set the decision threshold for classifying patients as responders versus non-responders.

**Supplemental Figure 7.**
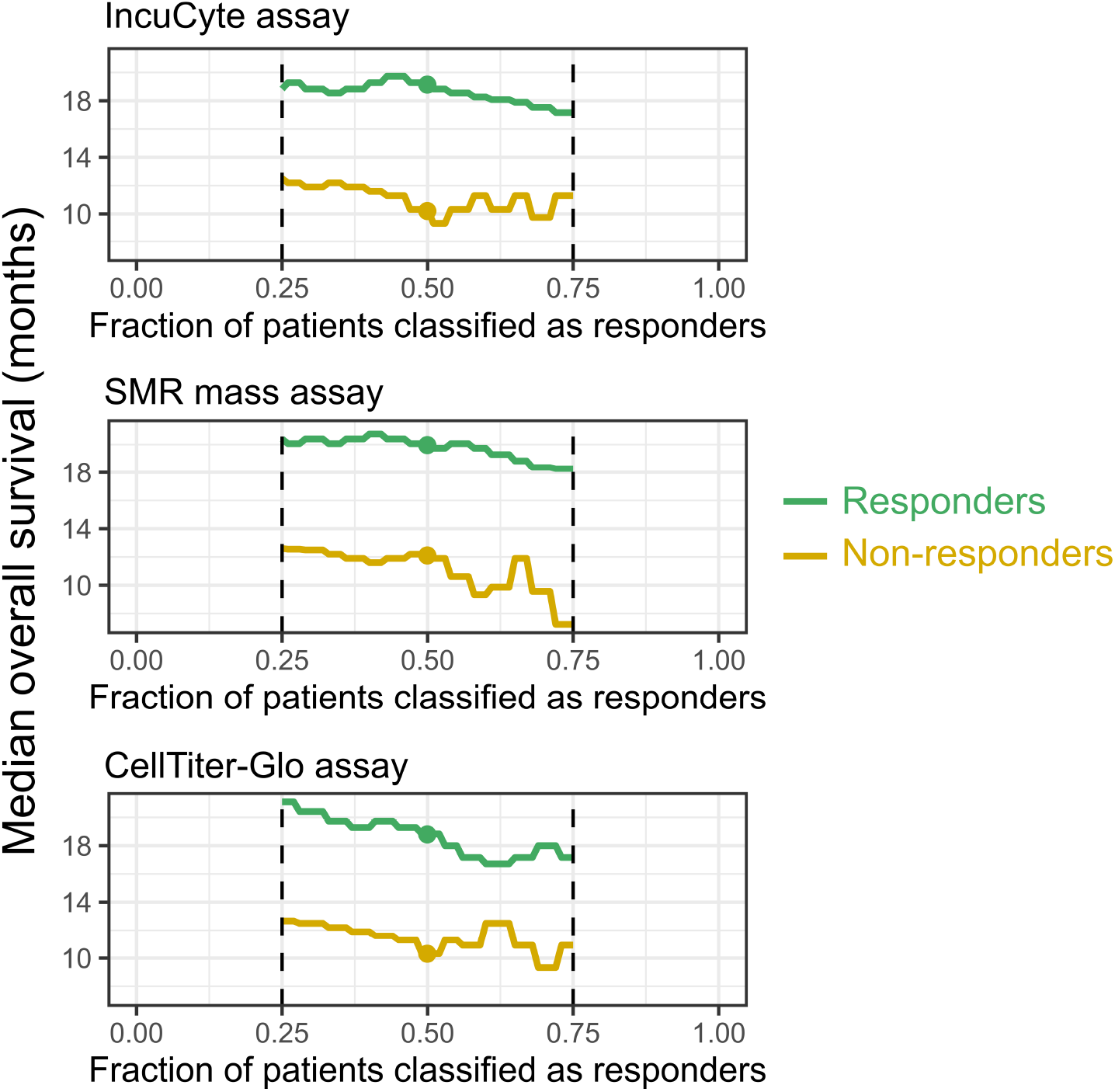
Predictive power depends only weakly on the fraction of patients classified as responders. We chose to classify the top 50% of most-responsive patients as “responders” for each functional assay, because approximately half of GBM patients respond to TMZ. Changing this number (here, shown for the range of 25% to 75% of patients classified as responders) only weakly affects the predictive power of the functional assays, i.e., the difference in survival between responders and non-responders.

**Supplemental Figure 8.**
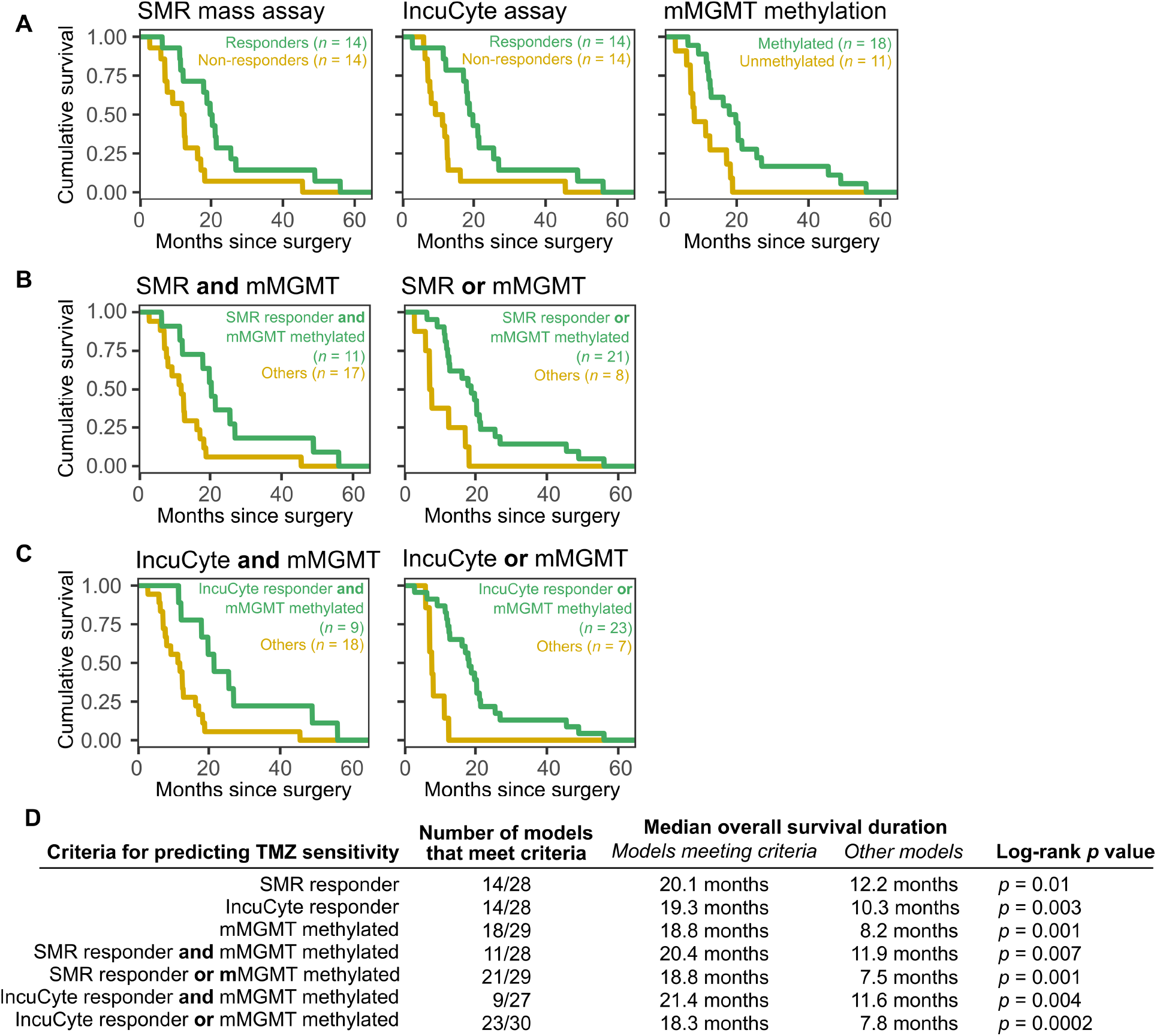
Combining functional and genomic biomarkers to predict patient outcome. (**A)** Survival distributions for patients stratified by single biomarkers, as shown in Fig. 6C. **(B)** Survival distributions for patients stratified by the combination of the SMR functional biomarker and the mMGMT methylation biomarker. **(C)** Survival distributions for patients stratified by the combination of the IncuCyte functional biomarker and the mMGMT methylation biomarker. **(D)** Summary statistics comparing the predictive power of combining functional biomarkers with MGMT to predict TMZ sensitivity. Depending on the criteria used to classify patients as likely TMZ-sensitive, one can achieve the desired level of sensitivity and specificity for predicting survival duration.

